# A pooled image-based CRISPR screen identifies EAF1 as a *T. gondii* modulator of ESCRT subversion

**DOI:** 10.64898/2026.07.08.737057

**Authors:** Einar B. Olafsson, Christophe-Sebastien Arnold, Jacob A. Kellermeier, Patrick A. Rimple, Hargobinder Kaur, Yifan Wang, Jonathan Z. Sexton, Staffan Svärd, Vern B. Carruthers, Matthew J. O’Meara

## Abstract

Intracellular pathogens remodel host cells by redirecting cellular machinery to the host-pathogen interface. The protozoan parasite *Toxoplasma gondii* co-opts host ESCRT proteins at the parasitophorous vacuole membrane (PVM), where they support budding of host-derived vesicles into the vacuole. Yet the parasite effectors that direct this process remain largely unknown. Here we developed spaCR (**s**patial **p**henotype **a**nalysis of **CR**ISPR-Cas9 screens), a pooled image-based screening framework that combines deep-learning classification of single-cell spatial phenotypes with well-level barcode genotyping and regression-based deconvolution of gene effects. Screening *T. gondii* secretory proteins recovered the known TSG101 recruiter GRA14 and identified an uncharacterized effector, ESCRT-association factor 1 (EAF1), that promoted TSG101 recruitment to the PVM. Targeted deletion confirmed its role across Type I and II lineages, and IP-MS recovered ESCRT-I and ESCRT-III components, including TSG101. Together, these findings identify EAF1 as an ESCRT-associated parasite effector and establish spaCR as a general framework for linking CRISPR perturbations to spatial phenotypes.

## 1 Introduction

*Toxoplasma gondii* (*T. gondii*) is an obligate intracellular parasite and one of the most tractable models for studying host-pathogen interactions^1^. From within its intracellular niche, the parasitophorous vacuole (PV), *T. gondii* orchestrates extensive remodelling of host membrane traffic^2^, including recruitment of the host endosomal sorting complex required for transport I (ESCRT-I) subunit tumour susceptibility gene 101 (TSG101) to the PV membrane (PVM)^3^. Recruitment of ESCRT to the PVM supports parasite nutrient acquisition by promoting ingestion of host cytosolic proteins and entrapment of Rab11A-positive, lipid-laden host vesicles within the PV^4^. Two PVM-resident parasite effectors are known to engage host ESCRT components. GRA64 (TGGT1_202620), immunoprecipitates with multiple ESCRT-I and accessory subunits^5^, and dense granule protein 14 (GRA14) binds the ubiquitin (Ub) E2 variant (UEV) domain of TSG101 through a cytosolically exposed Pro-Thr-Ala-Pro (PTAP) motif^3^. Recruitment of TSG101 to the PV is reduced but not abolished in GRA14-deficient parasites, indicating that additional effectors are involved^3^. However, fitness screening in a Δ*gra14*-sensitized background did not recover further ESCRT-engaging effectors^6^, highlighting a broader limitation of selection-based genetic screens where redundancy can mask function. In the case of ESCRT recruitment, redundancy may arise from both multiple parasite effectors that engage ESCRT and overlapping functions among ESCRT-I subunits, allowing loss of a single parasite factor to reduce TSG101 recruitment without causing a measurable fitness defect. Since recruitment of TSG101 to the PV is a spatial phenotype, we hypothesize that it can be better measured by imaging-based methods that resolve spatial features rather than readouts of parasite growth or indirectly by the abundance of transcripts or proteins.

Recent advances in robotic high-content microscopy and machine-learning-based image analysis have made morphological profiling a versatile platform for phenotypic screening, provided high-quality controls are available to train classifiers. However, because of the morphological heterogeneity of spatial phenotypes and the challenge of defining generalisable representations, it remains difficult to infer causal mechanisms from purely observational morphological profiling experiments. Pooled CRISPR-Cas9 perturbation screens have emerged as a scalable approach for establishing causal links between genotype and phenotype^7,8^. Yet they typically rely on bulk readouts such as mutant abundance shifts after sorting^9,10^ or growth selection^11–15^. Arrayed screens support measuring spatial phenotypes through imaging readouts but generating and maintaining one mutant per well limits throughput and inflates costs. Inherently, both formats are limited by low information density. Pooled screens constrain output complexity, while arrayed screens restrict input complexity.

Recent methods address this information-density trade-off by linking perturbations to genetically encoded barcodes decoded at single-cell resolution. Perturb-seq^16^ connects perturbations to transcriptional states by single-cell RNA sequencing, but does not directly capture spatial phenotypes such as protein localization or host-pathogen interface architecture. CRISPRmap^17^ and PerturbView^18^ extend this logic to spatial or multimodal readouts, but require specialized *in situ* decoding or sequencing workflows. Microraft arrays preserve image phenotype by isolating imaged clonal populations before barcode recovery^19^, and *in situ* genotyping of pooled strain libraries decodes barcodes inside imaged colonies^20^, but these formats are less readily adapted to dispersed single-cell infection phenotypes. Optical pooled screening^21^ most directly combines image-based phenotyping with single-cell perturbation identity, but requires multi-cycle *in situ* amplification and sequencing chemistry in the imaged sample. Together, these methods solve single-cell perturbation assignment, but at the cost of specialized infrastructure, terminal barcode decoding or assay formats that constrain the range of image-based phenotypes that can be measured (**Supplementary Table 1**).

A complementary strategy for increasing information density is to multiplex CRISPR mutation into each cell, and computationally deconvolve the causal associations between perturbation and cellular phenotypes^22^. When this strategy is paired with microscopy, the phenotype is read directly from images rather than from sequencing, so deconvolution depends on classifying the phenotypic state of single cells. Training accurate single-cell phenotype classifiers, however, requires carefully assembled datasets, and existing tools do not streamline the path from raw images to annotated training sets, creating a barrier for this approach.

We hypothesize that tracking genetic perturbations at the single-cell level is not required to identify gene-level effects. If a pooled mutant population is distributed across the wells of a standard microtiter plate, the set of mutations in each well can be recovered by low-cost well-level barcoding, direct PCR, and short-read sequencing. The well-averaged phenotype scores derived from single-cell images can then be regressed against this well-level genotype distribution. In principle, if the set of genotypes that exhibit a phenotype are sparse and only a small number are randomly distributed into each well, it should be possible to identify the causal genotypes with multiple linear regression (MLR). To test this, we developed spaCR (spatial phenotype analysis of CRISPR-Cas9 screens), a method that links well-averaged image phenotypes to well-level genotypes by MLR to recover gene-level effects. The accompanying open-source Python library streamlines the assembly of annotated training sets, applies deep learning to classify spatial phenotypes, maps barcoded reads to well-level genotypes, and resolves gene-level effects by MLR. To support adoption, we implemented a graphical user interface for spaCR, and to support experimental design, we developed spaCRPower, an R package that simulates spaCR screens.

Here we apply spaCR to a pooled gRNA library targeting *T. gondii* secretory proteins^15^ to identify additional regulators of TSG101 recruitment to the PVM. We classify single-cell phenotypes in parallel with two complementary approaches, the gradient-boosted decision tree algorithm XGBoost^23^ and the hybrid vision transformer MaxViT^24^. Independent validation of the screen hits confirmed one previously uncharacterized effector, which we name ESCRT-association factor 1 (EAF1, TGGT1_225160). Clean knockouts in Type I (RH) and Type II (Pru) lineages showed that EAF1 promotes TSG101 recruitment in both backgrounds, and a Δ*eaf1*Δ*gra14* double knockout phenocopied Δ*gra14*. Immunoprecipitation of EAF1-HA, followed by liquid chromatography-tandem mass spectrometry (IP-LC-MS/MS), recovered ESCRT-I and ESCRT-III subunits, accessory factors, and GRA14. Together, these data establish EAF1 as an ESCRT-associated parasite effector at the vacuolar interface, and demonstrate spaCR as a framework for resolving the genetic determinants of spatial phenotypes in pooled CRISPR-Cas9 screens.

## 2 Results

### 2.1 Modelling power and design of pooled spatial phenotype screens with spaCRPower

The spaCR experimental design distributes a pooled mutant population across *seeder* plates, replicates each well into matched *imaging* and *sequencing* plates, and uses MLR to relate well-level genotype distributions to single-cell phenotype scores (**Fig. 1a**). To understand how the power of the design to identify hits depends on the library complexity, classifier sensitivity, and well-level mutant distribution, we developed spaCRPower, an open-source R package. spaCRPower simulates each stage of spaCR screens and fits a Bayesian MLR model with a sparsity-inducing horseshoe prior to the simulated data (**Fig. 1b,c**). Hit-identification accuracy is quantified as the area under the receiver operating characteristic curve (AUROC) for ranking ground-truth hit genes by the posterior mean of their gene-level regression weights.

**Figure 1.**
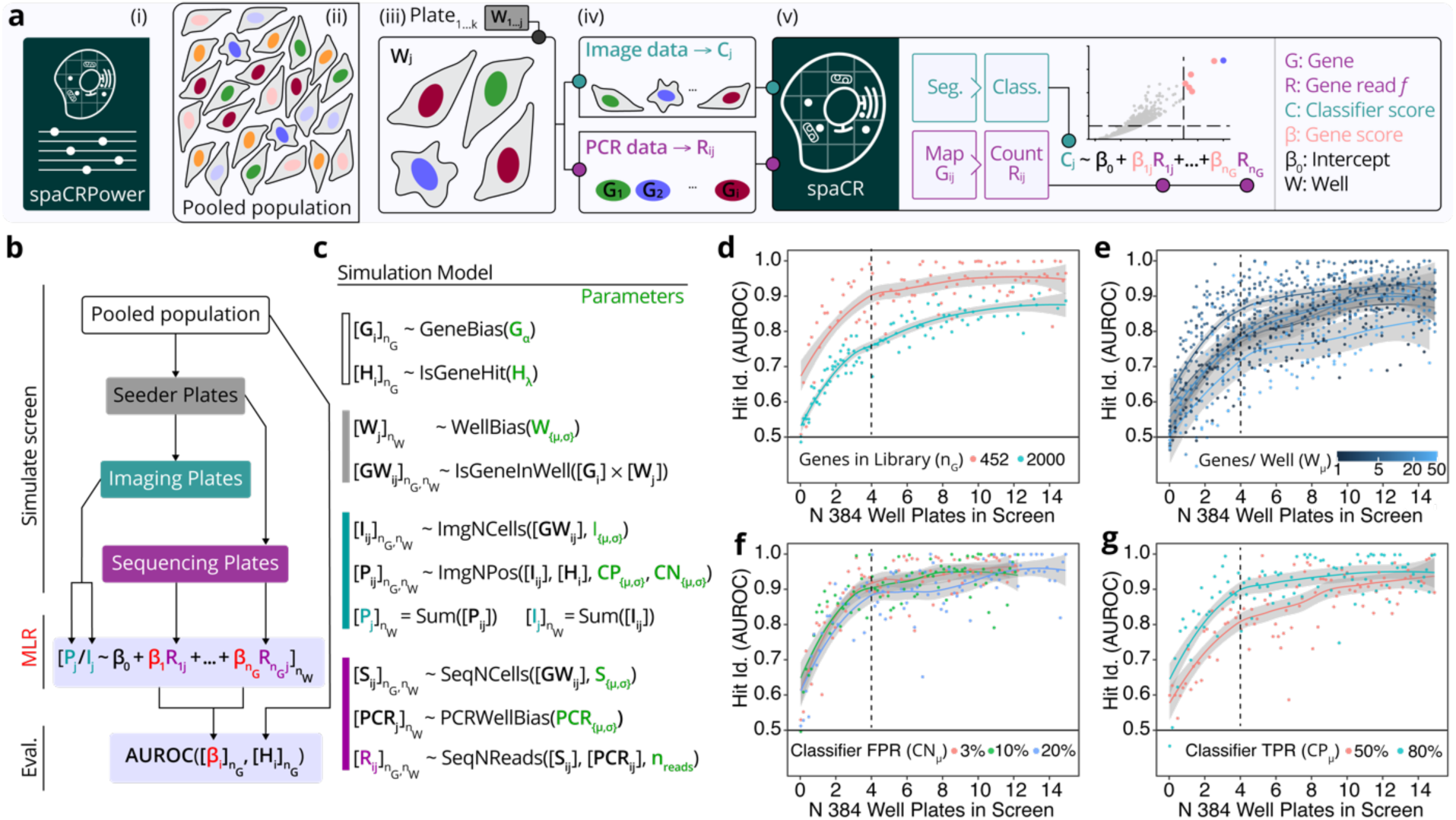
spaCR and spaCRPower workflows for pooled CRISPR-Cas9 imaging screens. ***a***, spaCR screen workflow. (i) spaCRPower estimates parameters for the screen design. (ii) A pooled population of cells carrying perturbations (G_1…i_) is distributed into (iii) plates (Plate_1…k_) with wells (W_1…j_). (iv) After proliferation, mutants are transferred to matched plates for genotyping (PCR followed by next-generation sequencing (NGS), FASTQ input to spaCR) and phenotyping (staining and imaging). (v) Within spaCR, images are segmented (Seg.) and classified (Class.) to yield well-level classification scores (C_j_). Barcoded reads are mapped (Map.) and counted to yield gRNA read fractions (R_ij_). C_j_ and R_ij_ are combined in multiple linear regression (MLR) to estimate gene-level effect sizes (β_i_). ***b***, schematic of the spaCRPower simulation workflow. ***c***, statistical simulation model summarized by stage. Parameter values are provided in Methods. ***d***-***g***, simulated hit-identification accuracy (AUROC, area under the receiver operating characteristic curve) as a function of the number of 384-well plates in the screen, assuming three 16-well control columns per plate. Parameters varied: ***d***, number of genes in the library (n_G_); ***e***, mean genes per well (W_μ_); ***f***, classifier false positive rate (FPR, CN_μ_); ***g***, classifier true positive rate (TPR, CP_μ_). Shaded bands indicate the 95% pointwise confidence interval for the mean prediction across 1,480 simulated screens. Dotted lines mark four plates, the design used in the TSG101 screen reported here.

To explore how the design parameters govern spaCR screen power, we used spaCRPower to simulate varying library complexity, well-level mutant distribution, and classifier accuracy. In the plate-number sweep, simulated screens of 452 genes reached AUROC > 0.9 with four 384-well plates, whereas larger screens at the scale of the *T. gondii* (∼8,000 genes) or *Homo sapiens* (∼20,000 genes) genomes required more than fourteen plates to reach the same accuracy (**Fig. 1d**). In the mutant-distribution sweep, the five-mutants-per-well on average design used below performed similarly to the simulated arrayed format (one-mutant-per-well), although with modestly lower AUROC (0.80 versus 0.85; **Fig. 1e**). Classifier false positive rates varied widely without affecting accuracy (**Fig. 1f**). However, reductions in the true positive rate caused accuracy to fall sharply (**Fig. 1g**), and phenotyping fewer than 100 cells per well dropped AUROC below 0.9 (**Extended Data Fig. 1g**). These results correspond to the experimental parameters used in the screen below. Fitting the simulation model to the four-plate TSG101 screen reproduced the empirical distributions of mutants per well, wells per gene, gene fractions per sequencing read, cells per well, and the proportion of positive cells per well (**Extended Data Fig. 1a-f**), and predicted AUROC = 0.90 (central 95% interval, 0.79–0.99; 99 simulations).

### 2.2 spaCR-enabled screening of TSG101 recruitment to the *T. gondii* PV

We previously showed that *T. gondii* recruits TSG101 to the PVM through the C-terminal PTAP motif of GRA14^3^. To identify additional parasite effectors involved in this process, we generated an RHCas9 strain expressing SAG1 signal peptide-DsRed^25^, which expresses DsRed in the PV lumen (RHCas9DsRed, **Extended Data Fig. 3a-b**) and transfected it with the host-parasite interactome gRNA library (Hi-Lib), a library containing 1,355 targeting gRNAs against 450 *T. gondii* genes (∼3 gRNAs/gene), together with 30 non-cutting control gRNAs. The targeted genes encode known and predicted^26^ GRA, rhoptry (ROP), and other signal-peptide-containing proteins^15^ (**Supplementary Table 2**). After drug selection, transfected parasites were distributed across four 384-well seeder plates at approximately five viable mutants per well and replicated for five days. Parasites were then transferred to matched imaging plates containing GFP-TSG101-tagged HeLa cells^27^ (HeLa^GFP-TSG101^) and sequencing plates for genotyping (**Fig. 2**). HeLa^GFP-TSG101^ cells infected with Δ*sag1* and Δ*gra14* (TGGT1_233460 gRNA 4 and TGGT1_239740 gRNA 1, **Supplementary Table 2**) parasites were used as negative (NC) and positive (PC) controls, respectively. The third column of each plate contained NC/PC mixtures at defined ratios as a holdout test set. The resulting workflow yielded paired well-level genotype distributions and single-cell phenotype data on which the regression analysis below was performed (**Extended Data Fig. 2)**.

**Figure 2.**
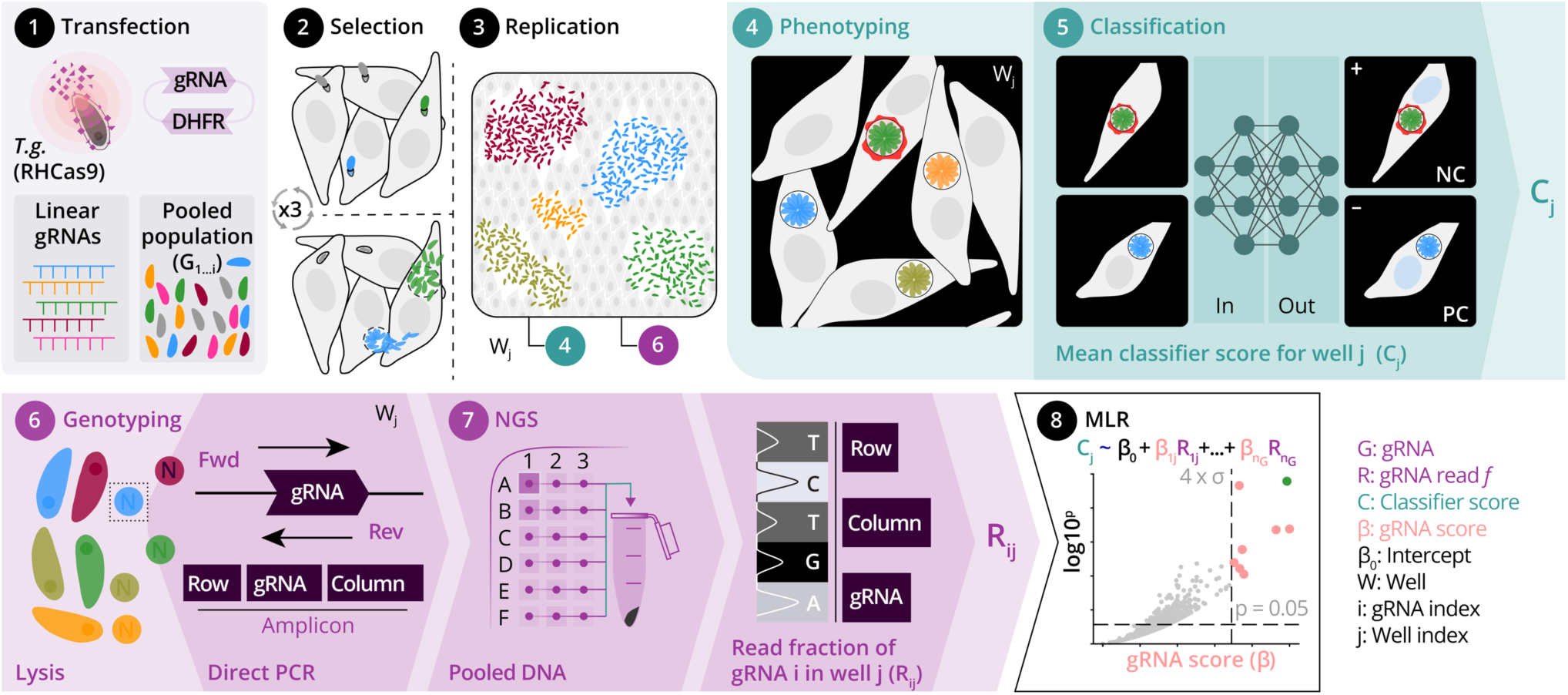
Proof-of-concept spaCR screen experimental workflow. (**1**) RHCas9DsRed *T. gondii* tachyzoites were electroporated with the Hi-Lib gRNA library and cultured on a human foreskin fibroblast (HFF) monolayer. (**2**) Drug selection over three passages and six days removed non-transfected parasites and parasites lacking the Cas9 or gRNA cassette. (**3**) Mutant parasites were distributed to confluent HFF monolayers in 384-well seeder plates at approximately five viable mutants per well and replicated for five days. (**4**) After plaque formation, parasites were transferred to matched imaging plates seeded with HeLa^GFP-TSG101^ cells, fixed at 24 h post-infection, stained, and imaged by confocal microscopy. Single-cell images were segmented for classification. (**5**) Images from negative-control (Δ*sag1*) and positive-control (Δ*gra14*) wells trained MaxViT and XGBoost classifiers, which were applied to cells from screening wells. (**6**) Parasites were also transferred to 384-well sequencing plates and subjected to direct PCR with row- and column-barcoded primers. (**7**) Pooled amplicons were sequenced by NGS and decoded to assign gRNAs to wells. (**8**) MLR estimated the effect size (β) of each genotype (R_ij_) on shifts in phenotype score (C_j_).

### 2.3 MaxViT outperforms feature-engineered XGBoost on mixed-ratio holdout data

Masks were generated for host cells, nuclei, and parasites (**Fig. 3a**), followed by extraction of measurements and single-cell images. As previously shown^3^, RHΔ*gra14* parasites (PC) recruited significantly less GFP-TSG101 than RHΔ*sag1* control parasites (NC) (**Fig. 3b**). To classify cells as PC or NC, we trained one MaxViT vision transformer^24^ per plate on cropped single-cell images (**Extended Data Fig. 3c-f**). We also trained 15 XGBoost models^23^ per plate on different categories of engineered intensity and morphology features. On the hold-out test set (a random sample of 10% of the NC and PC data), most XGBoost models and all MaxViT models exceeded 90% accuracy (**Fig. 3c**). The mixed-ratio holdout data, however, separated the two approaches. We compared the predicted PC fraction in the mixed-ratio holdout set across plates with the corresponding next-generation sequencing (NGS)-derived ground-truth fraction, using mean absolute error (MAE) relative to the NGS-derived ground-truth fraction as the performance metric. MaxViT achieved the lowest MAE, and XGBoost^GFP-TSG101^ (trained exclusively on GFP-TSG101 intensity features) achieved the next-lowest MAE (**Fig. 3d,e** and **Extended Data Fig. 3g-i**). Other XGBoost models, particularly those trained on morphology and Hoechst features, learned dataset-specific shortcuts visible only on the mixed holdout. SHapley Additive exPlanations (SHAP), model-derived feature importance and permutation-importance analyses of XGBoost^GFP-TSG101^ identified maximum and 0.95-quantile GFP-TSG101 intensity in and around the parasite as the most informative features (**Fig. 4a-d, Extended Data Fig. 4**), and saliency maps showed that MaxViT attended preferentially to the GFP-TSG101 and Hoechst channels (**Fig. 4e-g**). XGBoost^GFP-TSG101^ and MaxViT scores were carried forward to the screen analysis.

**Figure 3.**
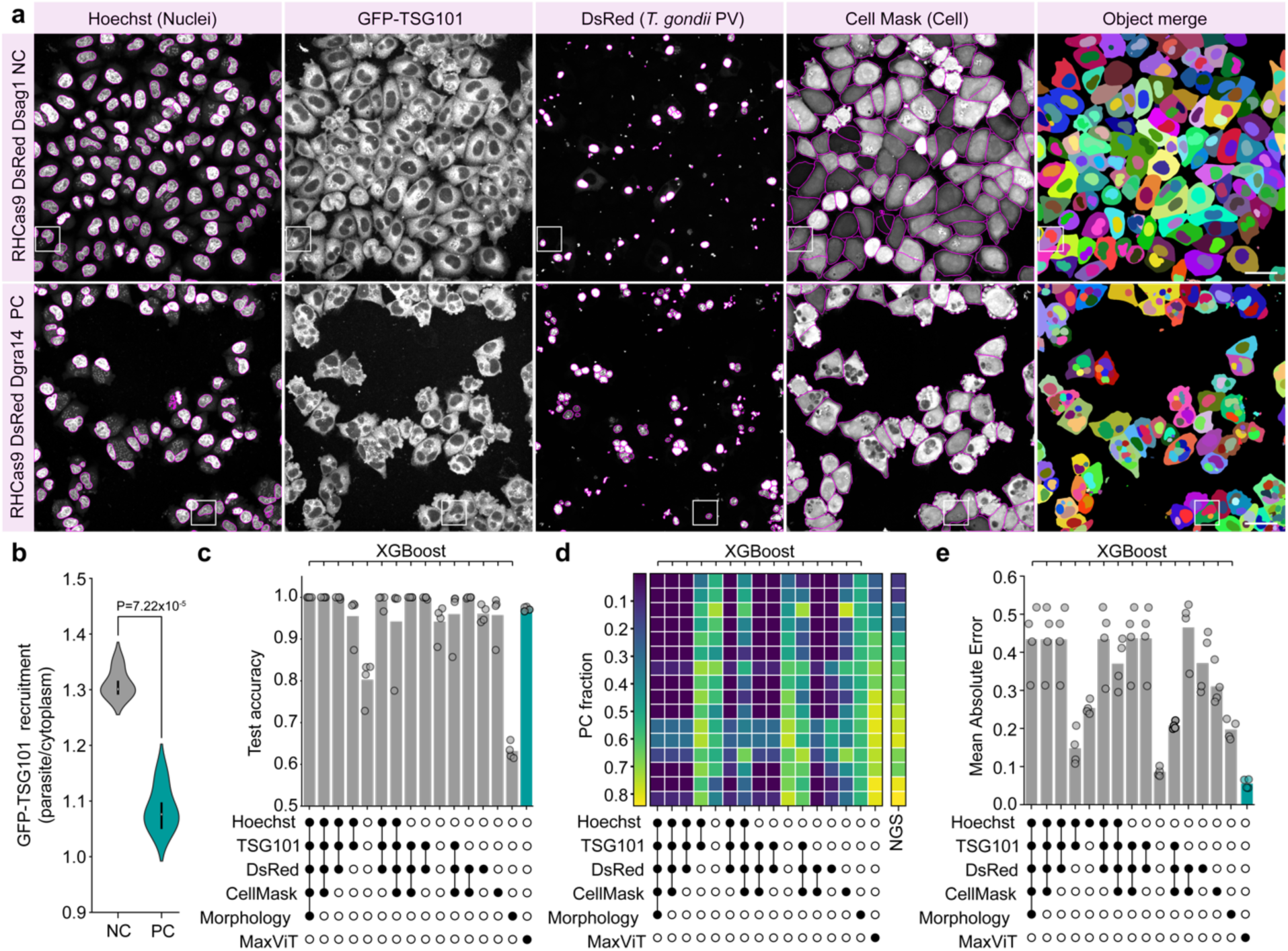
Single-cell phenotype classification of GFP-TSG101 recruitment. ***a***, representative confocal images of HeLa^GFP-TSG101^ cells infected with Δ*sag1* (negative control, NC) or Δ*gra14* (positive control, PC) RHCas9DsRed tachyzoites. Segmentation outlines (magenta) identify nuclei (Hoechst, first column), GFP-TSG101 (second column), parasites (DsRed, third column), and cells (CellMask, fourth column), and the merged segmentation across all object types (fifth column). White boxes indicate the cells highlighted in Fig. 4e,f. Scale bar, 100 µm. ***b***, well-average GFP-TSG101 recruitment in NC- and PC-infected wells (n = 4 plates, two-sided *t*-test). Violin plots show the full distribution. Horizontal lines indicate medians. ***c***, classification accuracy on the held-out test set (10% of data) for XGBoost models trained on the indicated feature combinations (grey bars) and for MaxViT models trained on cropped single-cell images (teal bar). One model was trained per plate per feature category (n = 4 plates). ***d***, performance on the mixed-ratio holdout (column 3 of each plate). NC and PC parasites were mixed at defined ratios. The resulting NC/PC fraction in each well was measured by NGS (right column, ground truth) and predicted by each classifier from (c). The heatmap shows PC fraction across rows for one representative plate (plate 1 of 4). ***e***, mean absolute error (MAE) between the NGS PC fraction and the classifier-predicted PC fraction for each model (n = 4 plates).

**Figure 4.**
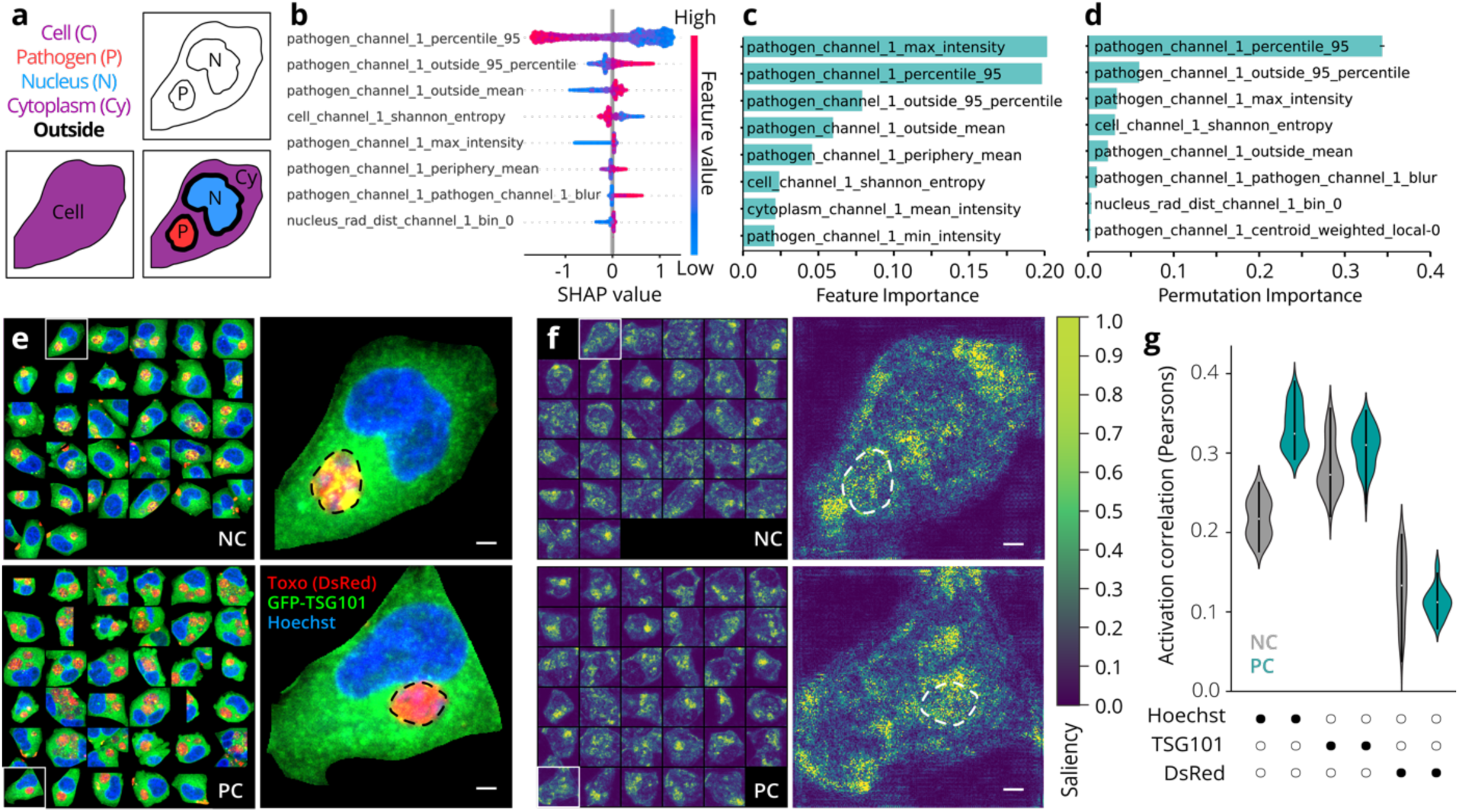
Feature attribution for XGBoost^GFP-TSG101^ and saliency analysis of MaxViT models. ***a***, schematic of the cellular compartments from which features and saliency are measured. Primary object: cell. Secondary objects: pathogen, nucleus. Tertiary objects: cytoplasm, nucleus periphery, pathogen periphery. Periphery objects are defined as the segmentation mask plus 5 px. ***b***, SHAP (Shapley additive explanation) feature importance, ***c***, model-derived feature importance, and ***d***, permutation importance for the eight top features of an XGBoost^GFP-TSG101^ model trained on PC and NC cells from plate 1 (representative of n = 4 plates). ***e***, representative single cells used to train MaxViT models (red, DsRed; green, GFP-TSG101; blue, Hoechst). Top, NC; bottom, PC. Scale bars, 5 µm. ***f***, saliency maps for the cells in (e), generated by one MaxViT model (representative of n = 4 models). Scale bars, 5 µm. ***g***, Pearson correlation between saliency-map activation and pixel intensity in each fluorescent channel (DsRed, GFP-TSG101, Hoechst), shown separately for NC and PC cells (n = 4 plates). Violin plots in (g) show the full distribution. Horizontal lines indicate medians.

### 2.4 Spatial phenotype CRISPR screen identifies EAF1 as a regulator of TSG101 recruitment

To estimate gene-level effect sizes, we combined per-well classifier scores, calculated by averaging cell-level scores within each well, with well-level genotype data from matched sequencing plates. We applied a 0.02 gRNA-fraction threshold per well for each gRNA, which yielded an average of five gRNAs per well, matching the distribution from the experimental design (**Extended Data Fig. 5a-d**). Wells with fewer than 100 phenotyped cells were excluded, leaving 587 wells (**Extended Data Fig. 5e-h**). Sequencing-quality controls confirmed that NC and PC reads were enriched in their assigned columns and that the mixed holdout series tracked with the expected gradient (**Extended Data Fig. 5i**). We averaged XGBoost^GFP-TSG101^ and MaxViT scores per well and fit MLR models with gRNA well-fractions as predictors. The screened gRNA library targeted predicted secretory proteins, including GRA, ROP and other signal-peptide-containing proteins (**Fig. 5a**). The most statistically significant hit using scores from both classifiers was TGGT1_239740_3, a gRNA targeting GRA14 (**Fig. 5b,c, Supplementary Tables 3-4**). Seven additional *T. gondii* genes satisfied our hit criteria in regressions on both XGBoost and MaxViT classifier scores (**Fig. 5b-c, Supplementary Tables 5-6**).

**Figure 5.**
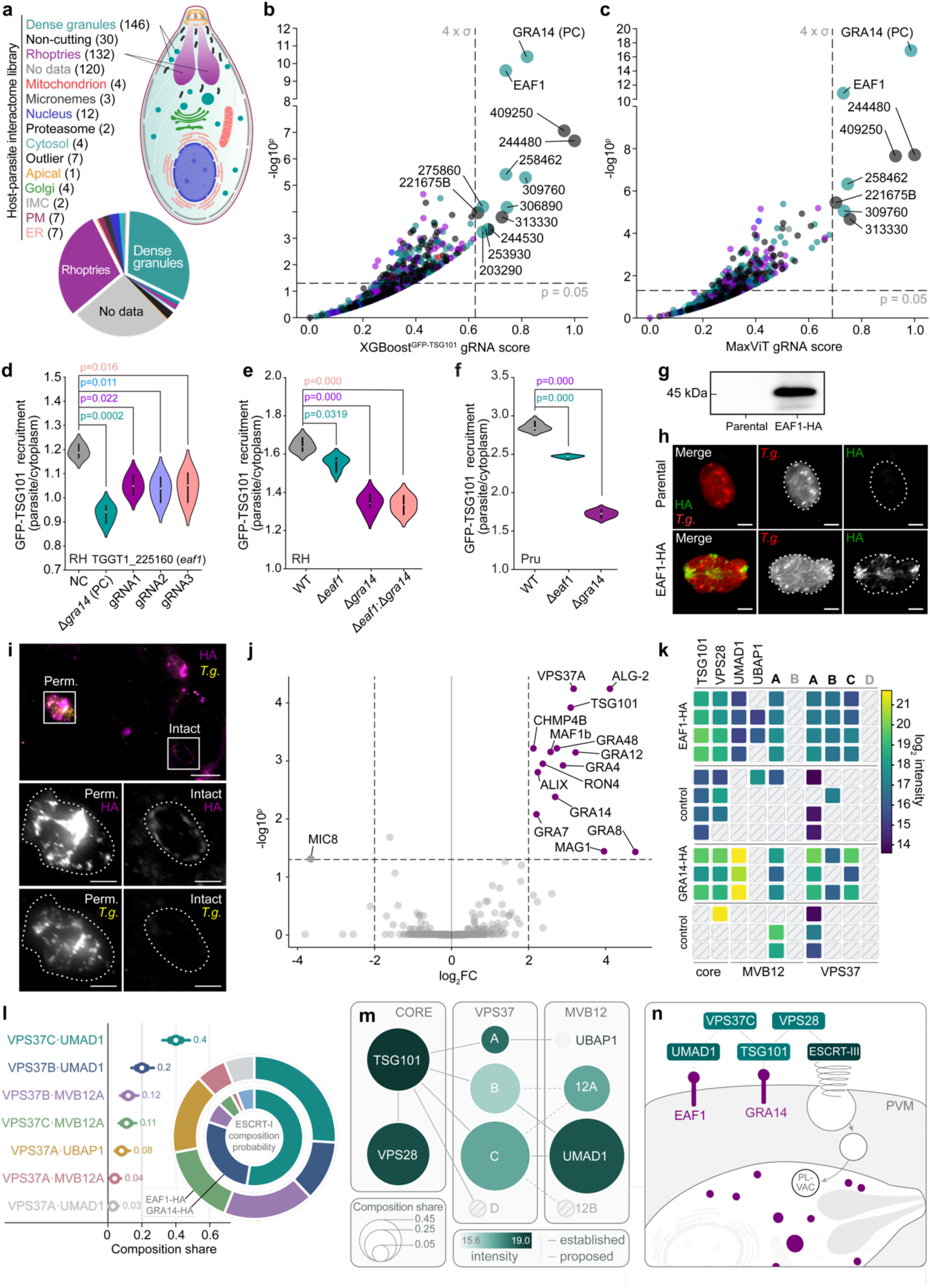
EAF1 promotes TSG101 recruitment to the *Toxoplasma* parasitophorous vacuole. ***a***, predicted subcellular distribution of the 450 genes in the host-parasite interactome library (Hi-Lib). Numbers indicate genes per category. ***b*, *c***, volcano plots of MLR results for well-level phenotype scores derived from XGBoost trained on GFP-TSG101 features (b, R^2^ = 0.699) or MaxViT (c, R^2^ = 0.715). Each point represents one gRNA. Colors indicate predicted compartment (legend in a). Dashed lines, p = 0.05 and 4 σ thresholds. Annotated genes pass both. ***d***, validation of EAF1 in RHCas9DsRed parasites transfected with individual gRNAs targeting TGGT1_225160 (gRNA1, gRNA2, gRNA3). GFP-TSG101 recruitment was quantified at 24 h post-infection (n = 3 independent experiments). ***e***, GFP-TSG101 recruitment in wild-type RH (RHΔ*hxg*), RHΔ*eaf1*, RHΔ*gra14*, and RHΔ*eaf1*Δ*gra14* parasites at 24 h post-infection (n = 4 independent experiments). ***f***, GFP-TSG101 recruitment in PruΔ*ku80* (Pru), PruΔ*eaf1*, and PruΔ*gra14* parasites (n = 4 independent experiments). For (d-f), one-way ANOVA with Tukey post-hoc test. Violin plots show the distribution and horizontal lines indicate medians. ***g***, anti-HA immunoblot of lysates from parental and EAF1-HA tagged parasites. Predicted molecular mass of EAF1-HA approximately 45.4 kDa (+3 kDa 3xHA). ***h***, anti-HA immunofluorescence in fibroblasts infected with parental (top) or EAF1-HA-tagged (bottom) parasites. HA signal is absent in the parental strain and localizes to the parasitophorous vacuole membrane in EAF1-HA. *T. gondii* (*T.g.*) is shown in red. Scale bar, 5 µm. ***i,*** selective permeabilization. HA signal is detected in PVM-intact vacuoles in which anti-Tg staining is absent, indicating access to the C-terminal HA epitope after host plasma membrane permeabilization. Anti-Tg staining marks fully permeabilized vacuoles. Scale bar, 5 µm. ***j***, anti-HA immunoprecipitation followed by liquid chromatography-tandem mass spectrometry (IP-LC-MS/MS) of EAF1-HA parasites compared with untagged control. Volcano plot highlights ESCRT-I, ESCRT-III, and ESCRT accessory proteins. Dashed lines, log_2_ fold-change ±2 enrichment and adjusted P = 0.05, n=4 biological replicates. **k,** Per-replicate log_2_ intensities of ESCRT-I subunits. Columns are the two invariant core subunits (TSG101 and VPS28), the four candidate MVB12 subunits, and the four VPS37 paralogs. Rows are the individual bait (EAF1-HA and GRA14HA) and untagged-control replicates. Color encodes the log_2_ intensity. Cells hatched in grey mark subunits not detected. **l,** Model-ranked inferred ESCRT-I compositions recovered with the two *T. gondii* effectors. The forest plot ranks the inferred subunit compositions by their combined model-derived share, with the median shown as a dot, the thick bar as the 50% credible interval and the thin bar as the 94% credible interval. The nested donut outer ring shows the EAF1 pull-down and the inner ring the GRA14 pull-down. ***m,*** ESCRT-I interaction network, with subunits arranged on assembly contacts. Edges mark documented subunit interactions (**Supplementary Table 16**). Node color is the mean bait log_2_ intensity of the GRA14 and EAF1 pull-down datasets. Node size indicates the model-derived composition share. ***n***, Working model in which EAF1 and GRA14 recover a VPS37C::UMAD1-enriched ESCRT-I population at the PVM. Lines and dotted lines indicate established and inferred interactions.

Next, we tested GFP-TSG101 recruitment of parasites transfected with gRNAs targeting screen hits. Parasites transfected with gRNAs targeting TGGT1_225160 recruited significantly less GFP-TSG101 than the Δ*sag1* control (**Fig. 5d**). We therefore named TGGT1_225160 ESCRT-association factor 1 (EAF1). Of the screen-hit candidates we tested, only TGGT1_225160 validated as a regulator of GFP-TSG101 recruitment, the other tested candidates did not validate as independent regulators (**Extended Data Fig. 5j-n**).

To distinguish stochastic false positives from systematic deconvolution artefacts, we examined how the candidate gRNAs were distributed across hit-positive wells. For most of the non-validated candidates we tested, the top-scoring gRNAs showed a well distribution correlated with either GRA14 gRNA 3 or EAF1 gRNA 2 (**Extended Data Fig. 5g**), suggesting enrichment through co-occurrence with true-positive gRNAs. Modelling effect-size misattribution with gene-level effects fixed at 1.0 for GRA14 and 0.7 for EAF1 reproduced the apparent significance of these false-positive gRNAs (**Extended Data Fig. 5h**). The screen therefore recovered two validated *T. gondii* regulators of TSG101 recruitment, with an observed false-positive rate within the range predicted by spaCRPower across plausible base hit rates for the library.

### 2.5 EAF1 promotes TSG101 recruitment in Type I and Type II *T. gondii* and acts on a shared pathway with GRA14

To confirm the role of EAF1 in a clean genetic background, we replaced the endogenous *eaf1* coding sequence with the HXGPRT cassette in RHΔ*hxg* and RHΔ*gra14* backgrounds (**Extended Data Fig. 6a**). RHΔ*eaf1* parasites recruited significantly less GFP-TSG101 than wild-type RHΔ*hxg* (P = 0.0319), and more than RHΔ*gra14*, consistent with **Fig. 5d**. The double knockout RHΔ*eaf1*Δ*gra14* phenocopied RHΔ*gra14* without further reduction (**Fig. 5e**), consistent with *gra14* being epistatic to *eaf1* and with the two effectors acting on a shared ESCRT-recruitment process rather than on independent additive pathways. The contribution of EAF1 to TSG101 recruitment was conserved across *T. gondii* lineages, with PruΔ*eaf1* parasites recruiting significantly less GFP-TSG101 than parental PruΔ*ku80* and showing an intermediate phenotype between PruΔ*ku80* and PruΔ*gra14* (**Fig. 5f, Extended Data Fig. 6b**).

### 2.6 EAF1 is a PVM-resident effector with a host-cytosol-facing C-terminus

Orthologues of EAF1 are confined to the Toxoplasmatinae (*T. gondii*, *H. hammondi*, *N. caninum* and *B. besnoiti*), with none recovered from other coccidia. Within *T. gondii*, EAF1 belongs to a paralogous family that also contains the GRA proteins GRA35 (TGGT1_226380), GRA36 (TGGT1_213067) and TGGT1_257970. The four orthologues formed a single well-supported clade distinct from EAF1 paralogs, consistent with duplications that preceded speciation (**Extended Data Fig. 7a, Supplementary Tables 7 and 8**). EAF1 comprises a predicted amino-terminal signal peptide (residues 20 to 43), a single transmembrane domain (residues 135 to 153) and a charged, low-complexity C-terminal domain (residues 154 to 401) with the properties of an intrinsically disordered region. Several short blocks in this domain are conserved across orthologues but not paralogs (**Extended Data Fig. 7b**). No canonical ESCRT-recruiting late-domain motifs were found in EAF1 (PT/SAP, PPxY or YPxL).

To confirm that EAF1 localizes to the PVM, we generated a strain expressing EAF1 endogenously tagged with three tandem C-terminal HA epitopes in a TgSS204 (RHΔ*ku80* DiCre) background (EAF1-HA, **Extended Data Fig. 6c**) and validated expression by anti-HA immunoblotting (**Fig. 5g**). HA staining was absent in the parental strain and localized to the PVM in EAF1-HA (**Fig. 5h**), confirming PVM localization of EAF1-HA.

To determine the membrane orientation of EAF1, we performed selective semi-permeabilization with saponin. An anti-*T. gondii* (α-*Tg*) antibody that stains the tachyzoite membrane was used to denote fully permeabilized infected cells (PV and host plasma membrane permeabilized, α-*Tg* positive) from those that were semi-permeabilized (host plasma membrane permeabilized, PV membrane intact, α-*Tg* negative). We found that EAF1-HA PVs in semi-permeabilized cells stained positive for anti-HA while α-*Tg* staining was absent, indicating that the C-terminal HA epitope is accessible from the host cytosol, placing the EAF1 C-terminus in the same compartment as the host ESCRT machinery (**Fig. 5i**).

### 2.7 EAF1 immunoprecipitation enriches host ESCRT-I components and GRA14

To identify EAF1 interactors, we performed IP-LC-MS/MS on lysates from EAF1-HA or control infected human foreskin fibroblasts (HFFs) using HA as bait. Bait recovery was confirmed by anti-HA immunoblotting (**Extended Data Fig. 6d**). Above the enrichment (log_2_ fold change > 2) and adjusted P < 0.05 thresholds, EAF1-HA significantly enriched the ESCRT-I subunits TSG101 (log_2_ fold change 3.09, adjusted P = 1.2 × 10^-4^) and VPS37A (3.17, 5.7 × 10^-5^). The ESCRT-III subunit CHMP4B (2.13, 6.1 × 10^-4^), the accessory factors ALIX (2.24, 1.6 × 10^-3^) and ALG-2 (4.11, 5.7 × 10^-5^), and GRA14 (2.69, 4.2 × 10^-3^) were also significantly enriched (**Fig. 5j-k, Supplementary Table 9**). At sub-FDR levels, recovery extended further to the ESCRT-I subunits VPS28 and MVB12A and the ESCRT-III subunit CHMP2A. The ESCRT-I subunits VPS37C and UMAD1 (UBAP1-MVB12-associated domain containing 1), the ESCRT-III subunits CHMP1A and CHMP3, and the ALG-2 partner peflin (PEF1) were detected exclusively or near-exclusively in EAF1-HA-infected samples (**Extended Data Fig. 8a**). By contrast, canonical ESCRT-0, ESCRT-II, and VPS4 components were not detected.

### 2.8 ESCRT-I composition recovered with EAF1 and GRA14

Because EAF1 and GRA14 converge on ESCRT-I, and recover multiple variable subunits whose identity determines complex function, we assessed which ESCRT-I species was most enriched by each bait. ESCRT-I comprises two constant subunits, TSG101 and VPS28, one of four VPS37 paralogues, VPS37A to VPS37D, and one MVB12-family subunit, UMAD1, UBAP1, MVB12A or MVB12B^28,29^. We scored the EAF1-HA and GRA14-HA^3^ IP-MS datasets with four established methods: SAINT^30^, SAINTexpress^31^, CompPASS^32^ and MiST^33,34^, (**Supplementary Tables 10 and 11)**. These methods ranked the variable subunits inconsistently (**Extended Data Fig. 8b,c**). MiST placed VPS37C and UMAD1 highest, whereas SAINTexpress favored the more abundant VPS37A, consistent with its abundance weighting. Moreover, because these methods score each prey independently, they do not account for the fixed architecture of ESCRT-I, in which VPS37 paralogues and MVB12-family proteins occupy alternative variable positions. We therefore applied a stoichiometry-constrained occupancy model (SCOM) to rank candidate VPS37-family and MVB12-family subunits within the defined ESCRT-I architecture, using bait enrichment and specificity against matched untagged controls in the EAF1-HA IP-MS dataset (IP-MS^EAF1-HA^, **Supplementary Table 9**), the GRA14-HA IP-MS dataset (IP-MS^GRA14-HA^)^3^ and the combined dataset (IP-MS^EAF1/GRA14^). For each variable position and inferred VPS37–MVB12-family pairing, SCOM reports the probability that a candidate has the largest relative share.

At the VPS37 position, the detected paralogs showed similar enrichment by bait-to-control comparison (IP-MS^EAF1-HA^ P = 0.13 and IP-MS^GRA14-HA^ P = 0.19, **Extended Data Fig. 8h**). VPS37A was recovered in most controls, consistent with its higher transcript^35^ and protein^36^ abundance relative to other VPS37 paralogues in fibroblasts (**Supplementary Table 12**). By contrast, VPS37C was not detected in any control sample (**Extended Data Fig. 8j**). SCOM ranked VPS37C as the leading VPS37-family subunit in both datasets, with the largest relative share in EAF1-HA and GRA14-HA immunoprecipitations (0.54 and 0.70, respectively; model-derived lead probability 0.99 and 0.98; **Extended Data Fig. 8d**).

At the MVB12-family position, UMAD1 was the most enriched subunit in IP-MS^GRA14-HA^ (P = 0.0002, **Extended Data Fig. 8i**). SCOM ranked UMAD1 as the leading MVB12-family subunit in IP-MS^GRA14-HA^ and IP-MS^EAF1/GRA14^ (model-derived lead probability 1.00; **Extended Data Fig. 8e**). In IP-MS^EAF1-HA^, the model assigned similar relative shares to MVB12A and UMAD1 (0.51 and 0.48, respectively; model-derived probability that UMAD1 leads = 0.39), and the two subunits could not be separated by enrichment alone (P = 0.21, **Extended Data Fig. 8i**). Thus, EAF1-HA recovered both MVB12A and UMAD1, whereas GRA14-HA preferentially recovered UMAD1 (**Fig. 5m, Extended Data Fig. 8f,g**). Under the independence assumption used to infer VPS37-MVB12-family pairings, VPS37C::UMAD1 was the top-ranked inferred pairing in IP-MS^GRA14-HA^ and IP-MS^EAF1/GRA14^ (shares 0.70 and 0.46, respectively; model-derived lead probability 0.98 and 0.99; **Fig. 5l, Supplementary Table 13**). In IP-MS^EAF1-HA^, VPS37C::UMAD1 and VPS37C::MVB12A had similar inferred pairing shares (0.26 and 0.27). Model-ranked compositions were robust to the Dirichlet concentration (**Extended Data Fig. 8k-m**) and to the imputation floor.

Together, these data are most consistent with EAF1 and GRA14 recovering a VPS37C::UMAD1-enriched ESCRT-I population at the PVM (**Fig. 5n**), with EAF1 also recovering MVB12A-containing ESCRT-I species.

## 3 Discussion

We developed spaCR, an open-source framework that links genetic perturbations to spatial single-cell phenotypes in pooled CRISPR-Cas9 screens by combining well-level genotyping, deep-learning image analysis and MLR. Applied to a screen of 450 *T. gondii* secretory proteins, spaCR recovered the known TSG101 recruiter GRA14 and identified EAF1 as a previously uncharacterized effector that promotes recruitment of host TSG101 to the PVM. Genetic, topological, and biochemical evidence place EAF1 alongside GRA14 as an ESCRT-associated parasite effector at the PVM, where the parasite co-opts the host ESCRT machinery.

With its C-terminal region exposed to the host cell cytosol, EAF1 is positioned to engage host machinery from the PVM. Deletion of EAF1 reduced TSG101 recruitment to the PVM, although the reduction was less pronounced than in Δ*gra14* parasites and the Δ*eaf1*Δ*gra14* double knockout did not further reduce TSG101 recruitment beyond the level observed in Δ*gra14* parasites. These data suggest that EAF1 contributes to the same ESCRT-recruitment pathway as GRA14 rather than acting through an independent additive pathway. While GRA14 uses a C-terminal PTAP motif to bind TSG101, we found no canonical ESCRT-binding motifs in EAF1. However, several short C-terminal regions were conserved among EAF1 Toxoplasmatinae orthologs, and to a lesser extent in EAF1 paralogs. Interestingly, one of the EAF1 paralogs, GRA35, recruits the HECT-type Ub ligase ITCH^37^. However, ITCH was not enriched in EAF1-HA immunoprecipitates, consistent with previous evidence indicating that EAF1 and GRA35 perform different functions^38^. The EAF1 interactome instead overlapped substantially with GRA64, which immunoprecipitates with several ESCRT proteins, including TSG101, VPS37A, VPS28, CHMP4B, and ALG-2^5^. This overlap is unlikely to reflect a direct interaction between EAF1 and GRA64, as neither GRA14^3^ nor EAF1 pulled down GRA64, and GRA64 was not recovered as a regulator of GFP-TSG101 recruitment in our screen. EAF1 also recovered the ESCRT-III subunits CHMP4B, CHMP2A, CHMP1A, and CHMP3, with no detectable recovery of ESCRT-0, ESCRT-II, or VPS4 components. Several PVM resident *T. gondii* proteins (GRA4, GRA7, GRA8, GRA12, GRA48, MAG1, MAF1b, and RON4) were co-enriched with EAF1, suggesting a wider effector platform.

To infer which ESCRT-I species were preferentially recovered with EAF1 and GRA14, we applied a stoichiometry-constrained compositional scoring model to the available IP-MS datasets. This analysis ranked VPS37C::UMAD1 as the leading inferred ESCRT-I composition in the combined EAF1/GRA14 dataset, while VPS37C::MVB12A was also supported in the EAF1 dataset. This distinction is biologically relevant because ESCRT-I complexes are compositionally specialized. UBAP1 preferentially assembles with VPS37A and defines an endosome-associated ESCRT-I complex involved in Ub-dependent multi-vesicular body (MVB) cargo sorting^29,39,40^. By contrast, UMAD1 preferentially associates with VPS37C or VPS37B and has been linked to cytokinetic abscission, a cargo-independent membrane-remodeling process^41^. MVB12A has been proposed to interact with VPS37C, but this pairing has not been experimentally validated^40,41^.

With VPS37 and UMAD1, the composition recruited to the PVM is more consistent with membrane-remodeling ESCRT-I than the canonical endosomal sorting pathway. This fits the topology of *T. gondii* ingestion, in which host-derived material is internalized across the PVM into the PV lumen^3,6^. ESCRT-dependent membrane scission operates in topologically related reactions, including intraluminal vesicle formation, viral budding and cytokinetic abscission^42,43^. Consistent with this, EAF1-HA recovered ESCRT-I and ESCRT-III subunits, but not ESCRT-0, ESCRT-II or VPS4 components.

The recovery of ALIX, ALG-2 and PEF1 further supports an ESCRT-associated membrane-remodeling environment at the PVM. ALIX links upstream ESCRT adaptors to ESCRT-III in cytokinetic abscission and retroviral budding^44,45^. ALG-2 is a Ca^2+^-binding adaptor that interacts with TSG101 and ALIX and can promote ALIX/ESCRT-III recruitment to damaged membranes independently of ESCRT-0 and ESCRT-II^46–48^. PEF1 forms heterodimers with ALG-2 and can modulate ALG-2-dependent trafficking functions^49,50^. Thus, EAF1-HA enriches a host ESCRT-associated protein set that includes ESCRT-I, ESCRT-III and Ca^2+^-responsive ESCRT accessory factors. Although these data define a set of ESCRT-associated proteins associated with EAF1, they do not resolve which proteins bind EAF1 directly. Because TSG101 and VPS28 are invariant ESCRT-I subunits, and the leading inferred EAF1-associated compositions differ mainly at the MVB12-family position, co-purification of VPS37C::UMAD1 and VPS37C::MVB12A is compatible with EAF1 associating with the ESCRT-I core, VPS37C, an MVB12-family subunit, ALIX/ALG-2-associated ESCRT machinery, or another PVM-associated factor. Defining the ESCRT-I species assembled at the PVM, and the ESCRT or ESCRT-associated factor that interacts directly with EAF1 should be a priority for future studies.

Fitness-based CRISPR screens in *T. gondii*, including screens performed in standard culture^12,14^ and *in vivo*^13,15^, have not identified EAF1, GRA14 or GRA64 as major fitness determinants. This may reflect partial redundancy among ESCRT-associated effectors, such that loss of one factor is buffered by others. Their contribution may also be context-dependent, as *T. gondii* infects most nucleated cell types across warm-blooded vertebrate hosts, whereas most functional data on ESCRT recruitment to the PVM have been generated in human and mouse cell lines. Roles for EAF1, GRA14 or GRA64 in other host species, cell types or life-cycle stages may therefore be missed by current fitness-screening conditions. Rather than measuring the fitness consequence of disrupting a pathway, spatial phenotype screens provide a more direct measurement that can be used to study the components of buffered systems. Synthetic lethal screens offer a complementary approach, pairing candidate knockouts with sensitizing perturbations to expose contributions that escape detection in isolation. Whether EAF1, GRA14, and GRA64 sustain ESCRT-I recruitment equivalently across life-cycle stages remains to be determined. Further, whether reduced TSG101 recruitment in Δ*eaf1* parasites produces an ingestion deficit similar to that reported for Δ*gra14*^3,6^ will require further study. Beyond these findings, several features of the screen warrant consideration as the approach is applied more broadly.

MLR treats each gRNA as an independent linear predictor and does not capture non-linear genetic interactions such as epistasis, synergy, threshold effects, or dose-response saturation. Where such interactions dominate, the regression step can be replaced by a hierarchical or non-linear model without altering the rest of the workflow. The estimates returned by this step depend on the quality of two inputs, the gRNA barcode assignments, and the well-level phenotype scores, both constrained by empirically determined thresholds. The 0.02 gRNA-fraction threshold retained the five gRNAs per well used in the screen and discarded low-fraction spurious barcodes and the 100-cell-per-well minimum marks the point above which well-level phenotype scores stabilized. Phenotype scores were produced by two classifiers, XGBoost^GFP-TSG101^ and MaxViT. On a mixed-ratio holdout, MaxViT generalized better than feature-engineered XGBoost models, several of which were accurate on standard test data yet had learned dataset-specific shortcuts. Supervised classification of this kind depends on labelled training data, obtained here from Δ*gra14* and Δ*sag1* controls. For phenotypes that lack equivalent controls, spaCR provides a manual annotation module in which single-cell images are labelled within the graphical user interface and stored in the project database for subsequent classifier training. Beyond these considerations, spaCRPower does not model cross-barcode sequencing errors, and Hi-Lib covers a 450-gene subset of the *T. gondii* secretome rather than the full ∼8,000-gene proteome, leaving parasite proteins upstream of GRA and ROP effectors outside the scope of the present screen. Genome-wide application is feasible with proportional scaling of plate number and sequencing depth, for which spaCRPower provides a quantitative basis.

Separating phenotyping from genotyping across paired imaging and sequencing plates offers practical advantages beyond the present screen. Because barcodes are not decoded in the imaged cells, imaging plates support non-destructive endpoints such as live-cell imaging and time-resolved phenotypes such as recruitment kinetics and organelle dynamics. These readouts are inaccessible to *in situ* sequencing and single-cell RNA sequencing, which require fixation or dissociation as terminal steps. Replicating one seeder plate into separate imaging plates also allows several spatial phenotypes to be scored in a single screen, each plate reporting a distinct phenotype.

In summary, this work establishes spaCR as a pooled CRISPR-Cas9 imaging framework for linking genetic perturbations to single-cell spatial phenotypes. Applied to *T. gondii* infection, spaCR recovered known regulators of TSG101 recruitment and identified EAF1 as an ESCRT-associated PVM effector, revealing an additional layer of parasite control over the host membrane-remodeling machinery. Because ESCRT pathways are exploited by diverse pathogens, these findings provide a framework for dissecting how intracellular pathogens engage conserved host membrane systems. More broadly, the spaCR workflow is not restricted to parasitic infection or fixed-cell endpoint assays. In principle, it is compatible with any proliferating cell type that can be transduced with a barcoded perturbation library, grown in microtiter plates, and analyzed by microscopy, provided that perturbation representation and phenotype scoring are sufficient.

## 4 Methods

### 4.1 Screen simulation (spaCRPower)

spaCRPower simulates each stage of a spaCR screen sequentially. Library: gene-level abundances G_i_ for i in [1, …, n_G_] are sampled from a Dirichlet distribution with shared concentration parameter G_α_ (alpha < 1 biases abundance toward a few genes), and gene-level hit status is sampled from a Bernoulli distribution with rate H_λ_. Seeder plates: per-well biases W_j_ for j in [1, …, n_W_] are sampled from a gamma distribution with mean W_μ_ and standard deviation W_σ_, and gene-well presence GW_ij_ is sampled from Bernoulli with probability G_i_ × W_j_. Imaging plates: cells per well I_j_ are sampled from a negative binomial distribution; per-gene cells in well j are drawn multinomially of size I_j_ over genes present; positive cells P_ij_ are sampled binomially of size I_ij_ with probability drawn from a beta distribution parameterized by classifier statistics CP(μ,σ) for hits and CN(μ,σ) for non-hits. Sequencing plates: per-gene cells S_ij_ are sampled from a negative binomial distribution; PCR amplification bias PCR_j_ is log-normal; reads R_ij_ are sampled jointly from a multivariate hypergeometric distribution over the population of all S_ij_ × PCR_j_. The model does not capture multi-guide knockout efficiency variation, cross-barcode sequencing errors, plate-position batch effects, or correlations between gene library abundance and phenotype.

Read counts per gene per well were normalized by the well-level total reads, offset by a 10^-4^ pseudo-count, and log-base-10 transformed. Wells with no imaged cells were excluded. To predict the number of positive cells per well, a Poisson generalized linear model with the log of total cells per well as offset was fit using BRMS^51^ in the Stan/R ecosystem (cmdstanr) with a sparsity-inducing horseshoe prior^52^ of 10 degrees of freedom over the gene-level coefficients β_i_, four independent NUTS chains of 8,000 samples each, max_treedepth = 12, otherwise default parameters. Hit-identification accuracy was quantified as AUROC for ranking ground-truth hit genes by the posterior mean of β_i_. For the *T. gondii* screen simulation: n_G_ = 452, G_α_ = 0.6, hit rate = 2.5%; n_W_ = 1,536, W_μ_ = 4.6; I_μ_ = 123, I_σ_ = √8000, CP_μ_ = 0.80, CP_σ_ = √0.1, CN_μ_ = 0.12, CN_σ_ = 0.01; S_μ_ = 1,000, S_σ_ = √1000, PCR_μ_ = 2, PCR_σ_ = 1, n_reads = 128,318. Parameter sweeps varied n_W_ from 10 to 5,010 in steps of 50, excluding non-converged runs. All parameters and scores are provided in the source data.

### 4.2 Parasite strain generation

To generate RHCas9DsRed, we first generated pDsRedHXGPRT by Gibson assembly (NEB E5510S) using primers 1-4 (**Supplementary Table 14**) bridging the chloramphenicol coding region of pDsRedCat^25^ and the HXGPRT cassette of pYFP.LIC.HXGPRT^53^. We then transfected 50 µg pDsRedHXGPRT into RHCas9 parasites and the DsRed-Sag1^1-188^-HXGPRT cassette was randomly integrated into the RHCas9 genome. Next, parasites underwent selection with mycophenolic acid (25 µg/mL) and xanthine (40 µg/mL) followed by limited dilution to generate monoclonal RHCas9DsRed. RHCas9DsRedΔ*gra14* and RHCas9DsRedΔ*sag1* were generated by Q5 mutagenesis (NEB E0554S) of pU6-DHFR-GRA14-gRNA1 (primers 5,6) and pU6-DHFR-SAG1-gRNA4 (primers 5,9), Sanger-verified, linearized with AseI, and transfected (50 µg) into RHCas9DsRed. Transfectants were selected in pyrimethamine (3 µM) and chloramphenicol (20 µM) and passaged three times before use.

RHΔ*eaf1*, RHΔ*eaf1*Δ*gra14*, and PruΔ*eaf1* strains were generated by replacing the endogenous *eaf1* coding sequence with the HXGPRT cassette using a Cas9-expressing plasmid encoding a single guide RNA targeting the N-terminus of *eaf1*. The Cas9-expressing plasmid was created using a Q5 Site-Directed Mutagenesis Kit (NEB; E0554S) and primers Q5MUT-EAF1-F and Q5MUT-EAF1-R (**Supplementary Table 14**). The HXGPRT cassette was amplified by PCR with homology arms corresponding to the 5′ and 3′ untranslated regions of *EAF1* using primers EAF1-conf-F and EAF1-conf-R (**See Supplementary Table 14**). Both constructs were co-electroporated into RHΔ*hxg*, RHΔ*gra14*^54^, and PruΔ*ku80* backgrounds. Transfectants were selected in mycophenolic acid (25 µg/mL) and xanthine (40 µg/mL) and cloned by serial dilution. Clones were screened by diagnostic PCR spanning the *EAF1* locus, with cassette insertion confirmed by an upward band shift relative to the parental line (**Extended Data Fig. 6a,b**).

EAF1-HA was generated in the TgSS204 (RHΔ*ku80* DiCre, gift of S. Lourido) background. The vector pG152-3xHA-HXGPRT-U1 was modified from pG152-LIC-HA-HXGPRT-U1 (gift of M. Meissner) by Q5 mutagenesis (primers 36, 37). The homologous region of *EAF1* was amplified by PCR (primers 38, 39) and cloned into the LIC site of pG152-3xHA-HXGPRT-U1 by PacI digestion and T4-polymerase-mediated annealing using dGTP (vector) or dCTP (insert), followed by transformation into NEB 5-alpha and Sanger verification. The construct was linearized with SphI and transfected into TgSS204. Transfectants were selected in mycophenolic acid (25 µg/mL) and xanthine (40 µg/mL) and cloned by serial dilution. HA-tagged EAF1 expression was confirmed by anti-HA immunoblotting and IFA (Fig. 5g, h).

### 4.3 Cell culture

Human foreskin fibroblasts (HFFs) and HeLa^GFP-TSG101^ cells^27^ were cultured at 37 °C and 5% CO_2_ in Dulbecco’s modified Eagle’s medium (DMEM) supplemented with 10% fetal bovine serum (FBS), 20 mM HEPES, penicillin/streptomycin (5 µg/mL), and 2 mM L-glutamine. *T. gondii* strains were maintained on HFF monolayers as previously described^6^.

### 4.4 Hi-Lib gRNA library transfection and partitioning

Hi-Lib (50 µg) was linearized with AseI, dialyzed, and electroporated (one 150 µs pulse at 1700 V) into 5 × 10^7^ RHCas9DsRed tachyzoites in cytomix (10 mM KPO_4_, 120 mM KCl, 5 mM MgCl_2_, 25 mM HEPES, 2 mM EDTA, 2 mM ATP, 5 mM glutathione, 0.15 mM CaCl_2_, 50 µg DNA, total volume 400 µL). Transfected parasites were transferred to a confluent HFF T75 monolayer. At 24 h post-transfection, medium was changed to selection medium containing pyrimethamine (3 µM), chloramphenicol (40 µM), and DNase I (10 µg/mL). Parasites were passaged three times under selection (6 × 10^6^, 3 × 10^6^, and 3 × 10^6^ parasites) before partitioning. Approximately 10 parasites were transferred per well across four 384-well plates seeded with confluent HFFs (∼5× library coverage), yielding approximately five plaques per well after five days. Mutants from the seeder plates were then replicated to matched HeLa^GFP-TSG101^ imaging plates and to 384-well sequencing plates.

### 4.5 Direct PCR and sequencing

Direct PCR was performed in 384-well plates (Bio-Rad HSP3805) with row- and column-barcoded forward and reverse primers (**Supplementary Table 15**) using the Phire Tissue Direct PCR Master Mix (Thermo Fisher F-170L). Briefly, 10 µL of dilution buffer with DNA release agent and 10 µL of sample from screen plates were mixed in matched 384-well plates, heated to 98 °C for 5 min, and 2 µL was transferred to direct-PCR plates containing 5 µL 2× master mix and 3 µL of forward and reverse primer (2 µM each), distributed as in **Supplementary Table 15**. PCR was run for 40 cycles. Pooled amplicons were concentrated by phenol/chloroform extraction (Sigma 77617-100ML), gel-purified (Qiagen 28604), and sequenced on a NovaSeq 6000 (GeneWiz), one plate per flow cell.

### 4.6 Immunofluorescent staining and imaging

HeLa^GFP-TSG101^ cells were seeded at 4,000 cells per well on poly-D-lysine-coated 384-well plates (Revity 6057302) 4h prior to challenge with *T. gondii*. After 24 h, samples were fixed (4% paraformaldehyde), stained with Hoechst (Sigma AMBH9A260690) and HCS CellMask Deep Red (Thermo Fisher H32721), and imaged on a Yokogawa CellVoyager 8000 dual spinning-disc confocal high-content microscope with a 40× water immersion objective and 50 µm pinhole. For each field of view (FOV), 10 z-slices were captured across 10 µm at 1 µm intervals. 20 FOVs were captured per well. For localization and semi permeabilization studies, images were acquired with a 40× objective on a Nikon Eclipse TI2 widefield microscope. All staining and storage buffers contained 0.04% sodium azide (Sigma 08591-1ML-F).

### 4.7 Selective permeabilization

To determine the membrane topology of EAF1, host plasma membrane was selectively permeabilized under conditions that leave the parasitophorous vacuole membrane (PVM) intact, allowing antibody access to host cytosol-facing epitopes but not to vacuolar lumen-facing or PVM-protected epitopes. Human foreskin fibroblasts (HFFs) were seeded on glass coverslips in 24-well plates and infected with EAF1-HA parasites at a multiplicity of infection of 1. At 24 h post-infection, samples were fixed in ice-cold 4% paraformaldehyde in PBS for 15 min on ice and washed three times with ice-cold PBS. Selective permeabilization was performed with 0.00001% (w/v) saponin in PBS for 5 min on ice, followed by three washes with ice-cold PBS. The saponin concentration was optimized by titration to leave some PVMs intact while permeabilizing the host plasma membrane and fully permeabilizing other vacuoles, confirmed by absence and presence of *T. gondii* staining. Low concentrations of saponin selectively extract cholesterol from the host plasma membrane^55^. After permeabilization, samples were blocked in 10% fetal bovine serum (FBS) in PBS for 30 min, incubated with the primary antibodies rat anti-HA (clone 3F10, Roche 11867423001) and anti-Toxo (Thermo Fisher PA1-7255) in 10% FBS in PBS for 1 h at room temperature, washed three times in PBS, and incubated with Alexa Fluor-conjugated secondary antibodies (Invitrogen) for 1 h at room temperature in the dark. Coverslips were mounted with ProLong Gold Antifade (Thermo Fisher P36930) and imaged on the Nikon Eclipse TI2 system as described in Section 4.6. PVM integrity under saponin treatment was confirmed by the absence of anti-Toxo staining. Images were processed identically across conditions in Fiji/ImageJ.

### 4.8 Immunoblotting

Lysates were resolved by SDS-PAGE, transferred to nitrocellulose, and blocked in 5% non-fat milk in TBST. Membranes were probed with anti-HA primary antibody (rat anti-HA, clone 3F10, Roche 11867423001) followed by HRP-conjugated secondary antibody. Signal was developed with SuperSignal West Pico PLUS Chemiluminescent Substrate (Thermo Fisher 34580) and imaged on a Syngene PXi6 system.

### 4.9 EAF1-HA immunoprecipitation and LC-MS/MS

HFFs in 15-cm dishes were infected with EAF1-HA or untagged TgSS204 parasites at MOI 3, four biological replicates per strain. At 24 h post-infection, monolayers were washed three times with cold PBS and scraped into cold lysis buffer (50 mM Tris-HCl pH 7.6, 200 mM NaCl, 1% Triton X-100, 0.5% CHAPS, complete protease inhibitor (Roche 11836153001)). Lysates were mechanically disrupted by passage through a 27.5G needle (5×) and sonicated on ice (1 s on, 1 s off, 20% amplitude, 30 cycles), incubated on ice for 30 min, and clarified at 1,000 × g for 10 min at 4 °C. Supernatants were incubated overnight with 0.25 mg anti-HA magnetic beads (100 µL slurry, Pierce 88836). Beads were washed once with lysis buffer and four times with wash buffer (50 mM Tris-HCl pH 7.6, 300 mM NaCl, 0.1% Triton X-100, complete protease inhibitor). The beads were washed twice with cold 100 mM ammonium hydrogen carbonate (AMBIC) and snap- frozen. The samples were processed at the IDEA National Resource for Quantitative Proteomics for LC-MS/MS analysis.

### 4.10 Image preprocessing and segmentation

Maximum intensity projections (MIPs) were generated from 10 z-slices spanning 10 µm per FOV. Filename metadata were extracted with the regular expression (?P<wellID>.*)_T(?P<timeID>.*)_F(?P<fieldID>.*)L(?P<laserID>.*)A(?P<AID>.*)Z(?P<sl iceID>.*)C(?P<chanID>.*).tif (specific to the Yokogawa CV8000). Channel images were combined into n-D arrays and segmented in Cellpose^56,57^: nuclei from channel [1] (cyto2_nuclei model, diameter 30, probability threshold 1), cells from channels [3,1] (cyto2 model, diameter 120, probability threshold 1), and *T. gondii* from channels [2,0] (cyto model, diameter 20, probability threshold 4). Channels were normalized before segmentation.

### 4.11 Single-cell extraction, classification, and regression

Fixed cells were stained, imaged, and segmented in spaCR using Cellpose^56,57^. For each segmented cell, intensity and morphological features were captured with scikit-image regionprops, supplemented by correlation, radial-distribution, and homogeneity measurements at the object perimeter and at perimeter +5 px. Cropped single-cell images were padded to 224×224 px and saved as plateID_wellID_fieldID_cellID.png. XGBoost^23^ models (spaCR.ml.ml_analysis) were trained on 90% of NC and PC measurements with 10% held out for testing, after removing non-numeric, low-variance, and highly correlated features. Permutation, feature, and SHAP importance were computed for each model. One MaxViT model^24^ was trained per plate to classify NC vs PC images using pretrained weights, focal loss, 8-fold image augmentation (flip, 90° rotation), and pixel normalization (mean [0.485, 0.456, 0.406], std [0.229, 0.224, 0.225]). Training used 100 epochs, batch size 64, gradient accumulation across four passes, AdamW optimizer (learning rate 10^-4^, weight decay 10^-5^, AMSGrad^58^) with reduce-on-plateau scheduling, 10% validation split, and dropout 0.1, on an NVIDIA RTX 3090 Ti (24 GB VRAM). PC and NC images were split into 80% training, 10% validation and 10% held-out test data, stratified by class.

Multiple linear regression was performed in statsmodels (v0.14.5) with the model C_j_ ∼ β_0_ + Σ_k_ β_k_^gRNA^ R_kj_, where C_j_ is the well-level phenotype score (mean MaxViT or median XGBoost) and R_kj_ is the read fraction of gRNA k in well j. A gene was classified as a hit if its regression p-value was < 0.05 and the absolute coefficient β exceeded the mean of non-targeting controls plus four standard deviations of the non-targeting distribution.

### 4.12 Multiple sequence alignment and phylogeny

Protein sequences of EAF1 (ToxoDB gene identifier TGGT1_225160) together with its orthologues and paralogs were retrieved from ToxoDB^59^ (release 68, VEuPathDB^60^). Orthologous and paralogous relationships were taken from the OrthoMCL groups recorded in ToxoDB. Within the recovered group, orthologues of EAF1 were present only in members of the Toxoplasmatinae, comprising *Toxoplasma gondii*, *Hammondia hammondi*, *Neospora caninum* and *Besnoitia besnoiti*. For the cross-species comparison one orthologue was used per species, namely *T. gondii* strain GT1 (TGGT1_225160), *H. hammondi* (HHA_225160), *N. caninum* (NCLIV_047520) and *B. besnoiti* (BESB_061290). Additional *T. gondii* strain models and duplicate annotation versions were effectively identical. The paralogs analyzed were the three in-paralogs encoded in the GT1 genome, namely TGGT1_213067 (GRA36), TGGT1_226380 (GRA35) and TGGT1_257970. Full-length protein sequences (**Supplementary Table 8**) were aligned with MAFFT (version 7) using the iterative local-pairwise mode (L-INS-i, options --localpair --maxiterate 1000)^61^. Approximate maximum-likelihood trees were inferred from the full-length alignment with FastTree^62^ (version 2.2) under the LG (Le and Gascuel) substitution model, with local support estimated from the Shimodaira-Hasegawa test. The signal peptide (residues 20 to 43) and the single transmembrane domain (residues 135 to 153) of EAF1 were set manually. The carboxy-terminal region to the transmembrane domain (residues 154 to 401), which is predicted to be exposed beyond the parasitophorous vacuole membrane (PVM), was treated as the C-terminal domain for the conservation analysis.

### 4.13 Conservation and motif analysis

Per-column amino acid identity was computed against the consensus of the four orthologues, with gaps excluded from the consensus and counted against identity. Conserved blocks were defined as runs of at least four consecutive alignment columns in which at least three of the four orthologues (75 % or more) shared the consensus residue. The C-terminal domain of EAF1 was screened for the three canonical late-domain motifs that recruit host ESCRT: PT/SAP (UEV of TSG101), PPxY (binds the WW domains of NEDD4-family HECT Ub ligases) and YPxL or LYPx(n)L (binds the V domain of ALIX), and for the minimal proline-directed phosphorylation consensus [S/T]-P, using regular expression.

### 4.14 Enrichment and specificity

Protein-level log_2_ intensities were taken for each subunit in the EAF1 IP-MS dataset (**Supplementary Table 9**) and the published GRA14 IP-MS dataset^3^. For the GRA14 dataset the peptide intensities were summed to the protein level for each replicate and log_2_ transformed. Missing values were treated as non-detection and, where a subunit was detected in at least one bait replicate, were imputed at a per-experiment floor for the enrichment calculation. Enrichment over the untagged control was estimated for each subunit as the difference of mean log_2_ intensities, with an empirical Bayes moderation of the residual variance across subunits to stabilize the estimate at the small replicate count^63^. The two experiments were combined by inverse-variance meta-analysis of the per-experiment enrichment estimates. Specificity was assessed with a Fisher exact test on the number of control replicates in which each subunit was detected, pooled across the seven control replicates of the two experiments. Subunits absent from the control were assigned maximal control specificity for the compositional scoring model.

### 4.15 Stoichiometry-constrained occupational modeling

The stoichiometry-constrained occupancy scoring model (SCOM) was designed in line with quantitative IP-MS and complex-centric proteomics approaches that use prey intensity, background correction and prior knowledge of complex architecture to infer interactor confidence, relative subunit stoichiometry or complex variants^64–66^.

SCOM ranks candidate ESCRT-I subunits within the two variable positions of the complex using the EAF1-HA and GRA14-HA^3^ immunoprecipitation datasets. VPS37A–D were treated as alternatives for the VPS37 position, and UMAD1, UBAP1, MVB12A and MVB12B were treated as alternatives for the MVB12-family position. For each subunit i, enrichment E_i_ was calculated as the difference in mean log_2_ intensity between tagged and untagged samples, with missing values imputed at a dataset-specific floor and negative enrichment values set to zero. Specificity S_i_ was defined as 1 − D_i_/n_C_, where D_i_ is the number of control replicates in which subunit i was detected and n_C_ is the total number of control replicates. Within each variable position p, candidate shares were calculated from enrichment weighted by specificity and normalized across all candidates in that position: Mi = E_i_ S_i_ / Σ_j∈p_ E_j_ S_j_. For each inferred VPS37–MVB12-family pairing of subunit a with subunit b, the pairing share was calculated as: C_ab_ = M_a_ M_b_. Pairing shares were renormalized across observed pairings. The two variable positions were treated as independent because the IP-MS data resolve subunit abundance but not direct co-membership within individual complexes. Relative position shares and inferred pairing shares were modelled using Dirichlet distributions, θ ∼ Dirichlet(κ_M_) for each variable position and θ ∼ Dirichlet(κ_C_) for inferred pairings, with concentration parameter κ set from the between-replicate dispersion of the shares. From N = 2 × 10^5^ posterior draws, the probability that a subunit or inferred pairing had the largest relative share, or exceeded another candidate, was calculated as the fraction of draws in which this condition was met. Positions, inferred pairings and pairwise comparisons were ranked by these probabilities.

SCOM does not infer direct co-membership of subunits within individual ESCRT-I complexes and does not model batch or platform differences between datasets, correlation between enrichment and specificity, or uncertainty in missing-value imputation beyond the reported floor-sensitivity analysis.

### 4.16 Statistical analysis

All hypothesis tests were two-sided at α = 0.05. Normality was tested with Shapiro-Wilk for small sample sizes (3 ≤ n < 8) or D’Agostino-Pearson K^2^ for larger sample sizes (n ≥ 8), and homoscedasticity with Levene’s test. Two-group comparisons used Student’s *t*-test (unpaired) or paired *t*-test for normal data and Mann-Whitney U or Wilcoxon signed-rank otherwise. Comparisons of more than two groups used one-way ANOVA (parametric) or Kruskal-Wallis (non-parametric), followed by Tukey HSD or Dunn post-hoc tests with Benjamini-Hochberg correction. Plots were produced with the seaborn v0.13.2 and Matplotlib v3.8.3 libraries in the spacr.plot module and curated in Adobe Illustrator 2026. Analyses used statsmodels v0.14.5, SciPy v1.12.0, and scikit-learn v1.4.1.

### 4.17 Data and code availability

spaCR (v1.3.4) is available at https://github.com/EinarOlafsson/spaCR and https://pypi.org/project/spacr/, with API documentation at https://einarolafsson.github.io/spacr/ and a step-by-step tutorial at https://einarolafsson.github.io/spacr/tutorial/. spaCRPower is available at https://github.com/maomlab/spaCRPower. Raw sequencing data are deposited in the NCBI BioProject (PRJNA1261935, SRA SRR33531217-SRR33531220). Image data are deposited at the EBI BioStudies repository (S-BIAD2135). The mass spectrometry proteomics data have been deposited to the ProteomeXchange Consortium via the PRIDE^67^ partner repository with the dataset identifier PXD080696. Intermediate datasets, training hyperparameters, and analysis parameters are provided as Supplementary Information.

## Extended Data figures

**Extended Data Figure 1.**
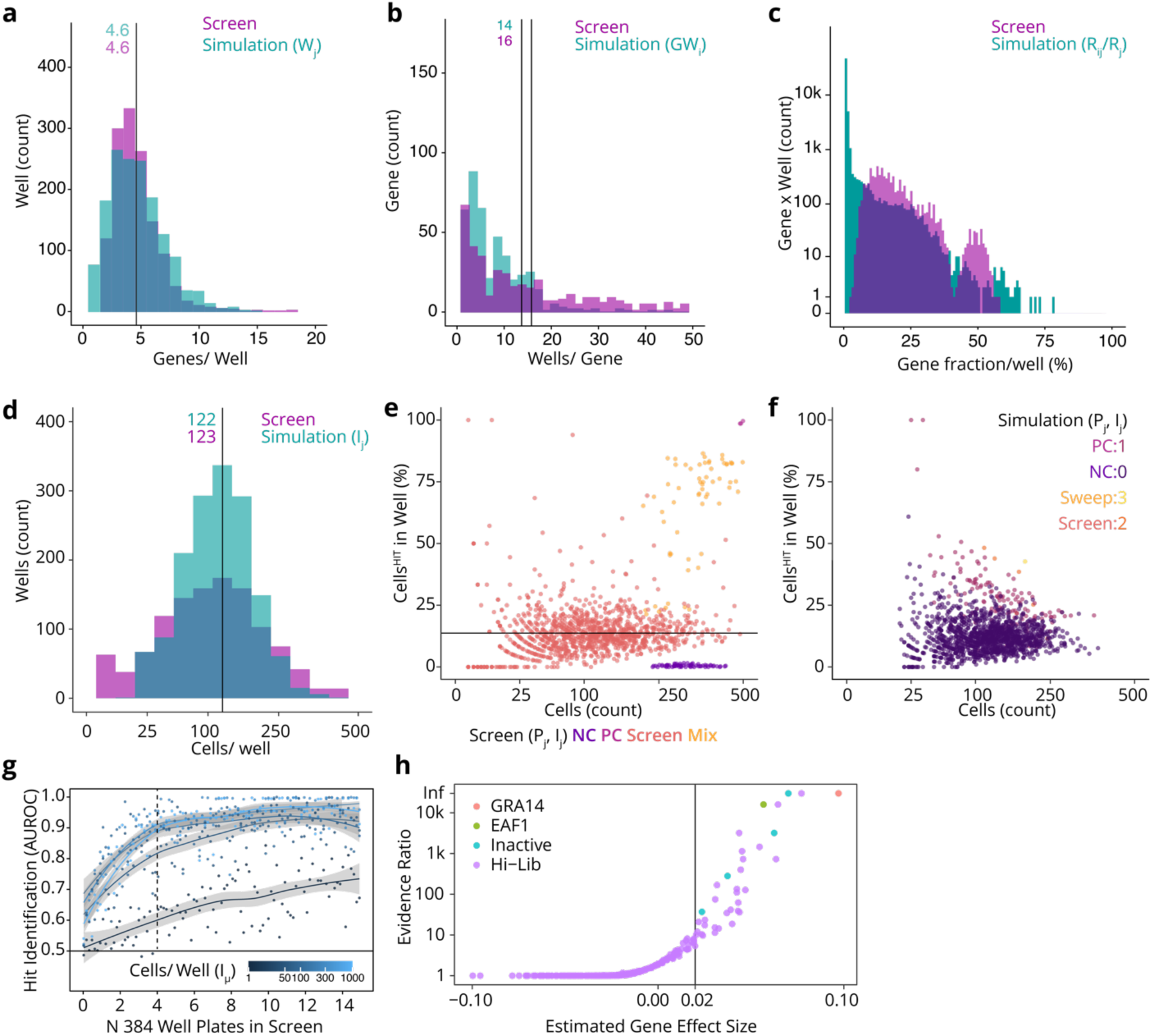
spaCRPower simulation parameters fitted to the TSG101 *T. gondii* screen. Comparison of summary statistics from spaCRPower simulations (parameters fitted to the empirical screen) with the corresponding statistics from the four-plate TSG101 screen reported in this work. Seeder plate: ***a***, histogram of gene counts per well; ***b***, histogram of well counts per gene. Sequencing plate: ***c***, histogram of per-gene reads as a fraction of total well reads. Imaging plate: ***d***, histogram of cells per well; ***e***, scatter plot of percent positive cells versus total cells per well in the screen, separated by control type (PC, NC, sweep) and screening wells; ***f***, same plot from simulated data, coloured by the number of positive genes in each well. ***g***, AUROC for hit identification as a function of the number of 384-well plates, with cells per well varied. Shaded band indicates the 95% pointwise confidence interval for the mean prediction across 989 simulated screens. Dotted line marks four plates as used in the TSG101 screen. ***h***, Bayesian MLR fit to the observed MaxViT classification and gRNA fraction data. The evidence ratio is the posterior probability that each β_i_ > 0.02 (the assumed noise threshold), and the gene effect size is the posterior mean of β_i_. The plotted genes pass the indicated posterior evidence threshold. EAF1 ranks among the top three by both effect size and evidence ratio.

**Extended Data Figure 2.**
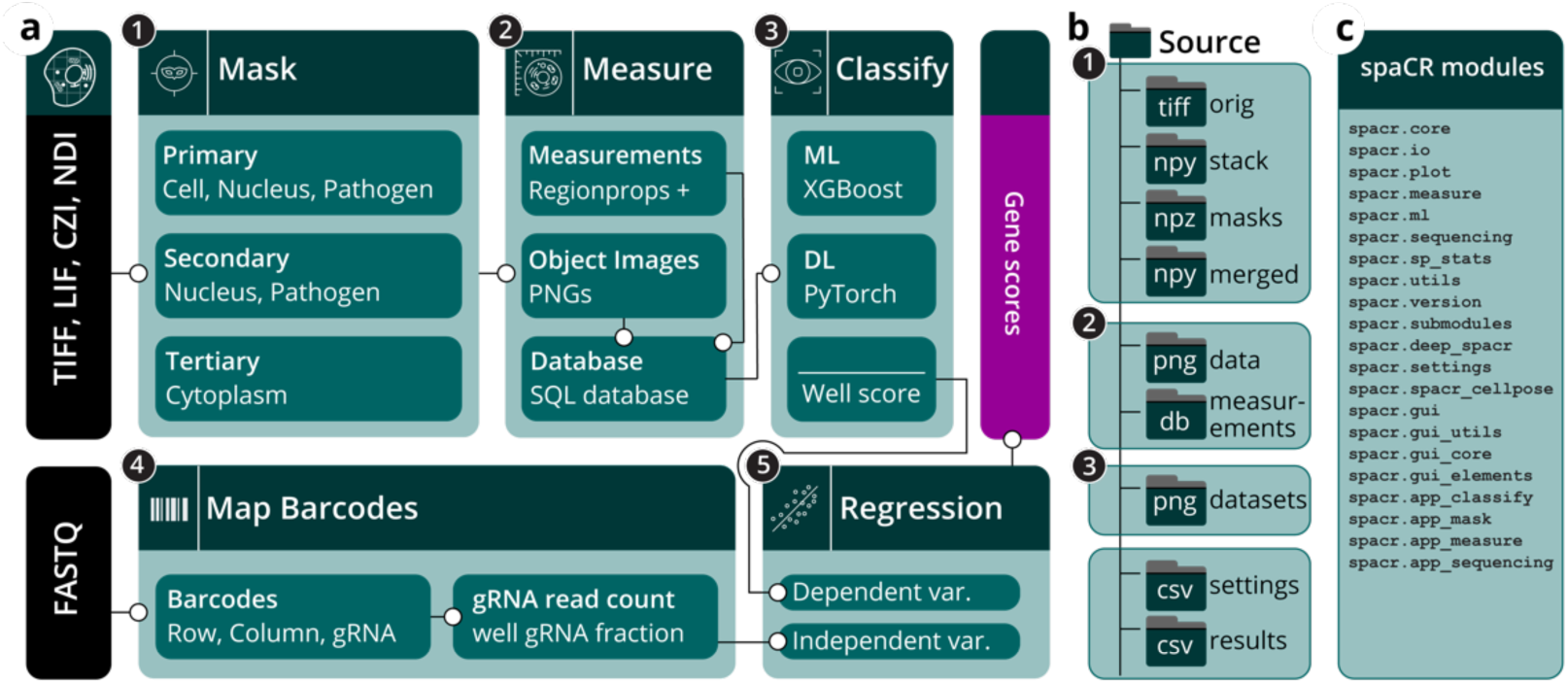
spaCR pipeline overview and data organization. ***a***, Schematic workflow of the spaCR pipeline for pooled image-based CRISPR screens. Inputs (black) are microscopy images (TIFF, LIF, CZI, NDI) and sequencing reads (FASTQ). Five graphical user interface modules (teal) handle the workflow: (1) Mask, generates object masks for cells, nuclei, pathogens, and cytoplasm; (2) Measure, extracts object-level features and crops single-cell images, storing data and image paths in a SQL database; (3) Classify, applies machine-learning (ML, XGBoost) or deep-learning (DL; PyTorch) models to classify objects and summarizes results as well-level scores; (4) Map Barcodes, maps row, column, and gRNA barcodes from sequencing reads to wells; (5) Regression, estimates gRNA effect sizes by MLRon well-level summary statistics. ***b***, spaCR output folder structure, including locations for raw and processed images, masks, object-level measurements, datasets, and analysis results. ***c***, spaCR Python library modules.

**Extended Data Figure 3.**
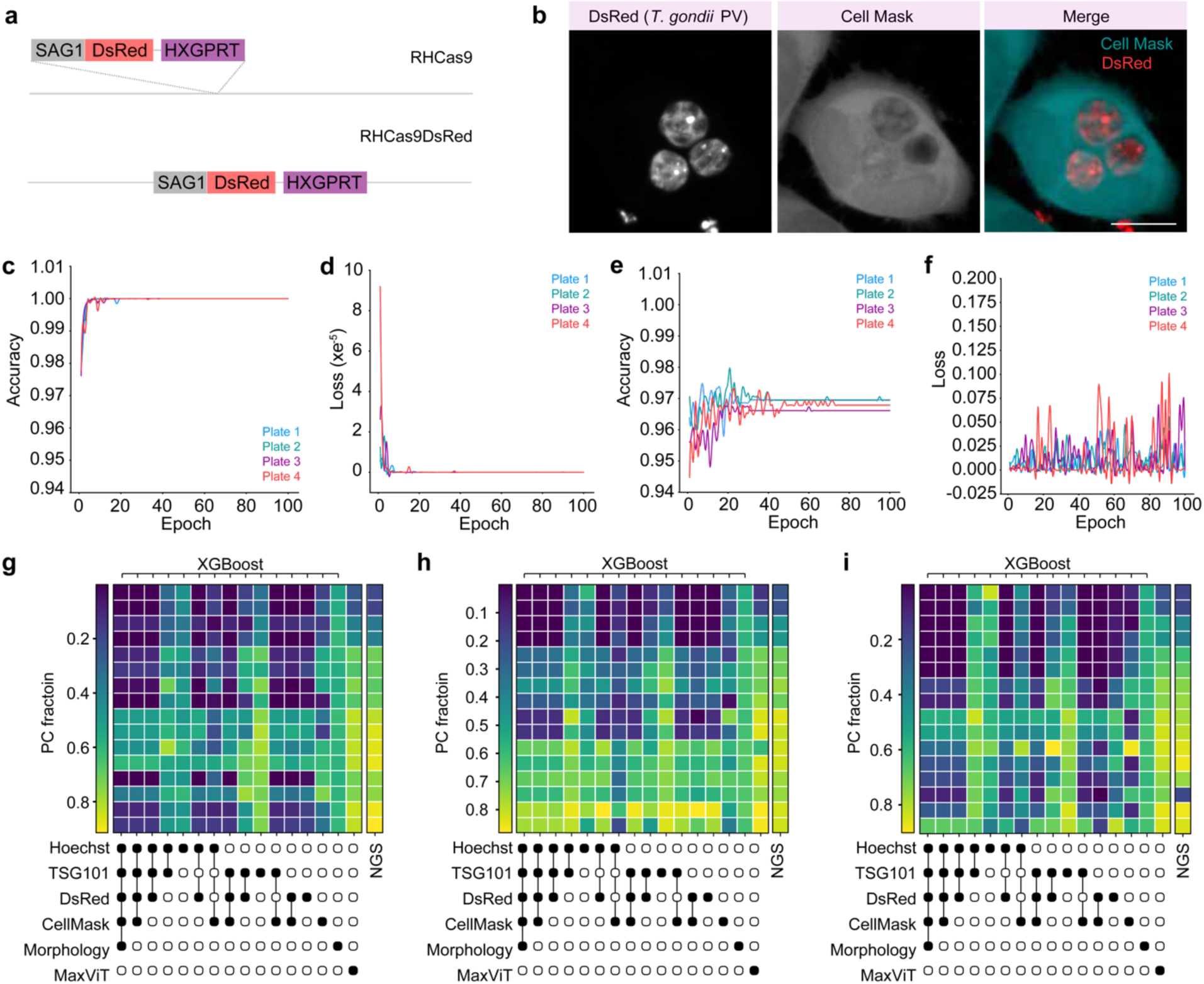
RHCas9DsRed generation and per-plate XGBoost and MaxViT validation. ***a***, schematic of the DsRed insertion strategy used to generate RHCas9DsRed (see Methods). ***b***, confocal images of HeLa cells infected with RHCas9DsRed parasites at 24 h post-infection, stained with CellMask. Left, luminal DsRed expression; middle, CellMask-stained host cell; right, merge. Scale bar, 10 µm. ***c***-***f***, training and validation curves over 100 epochs for one MaxViT model per plate (n = 4 plates). HeLa^GFP-TSG101^ cells infected with PC and NC parasites from columns 1 and 2 were used for training (80%), validation (10%), and testing (10%). ***c***, training accuracy; ***d***, training loss; ***e***, validation accuracy; ***f***, validation loss. ***g***-***i***, performance on the mixed-ratio holdout for plates 2-4 (compare with Fig. 3d, plate 1). RHCas9DsRedΔ*sag1* (NC) and RHCas9DsRedΔ*gra14* (PC) parasites were mixed at defined ratios. The NC/PC fraction was measured by NGS (right column, ground truth) and predicted by each classifier indicated.

**Extended Data Figure 4.**
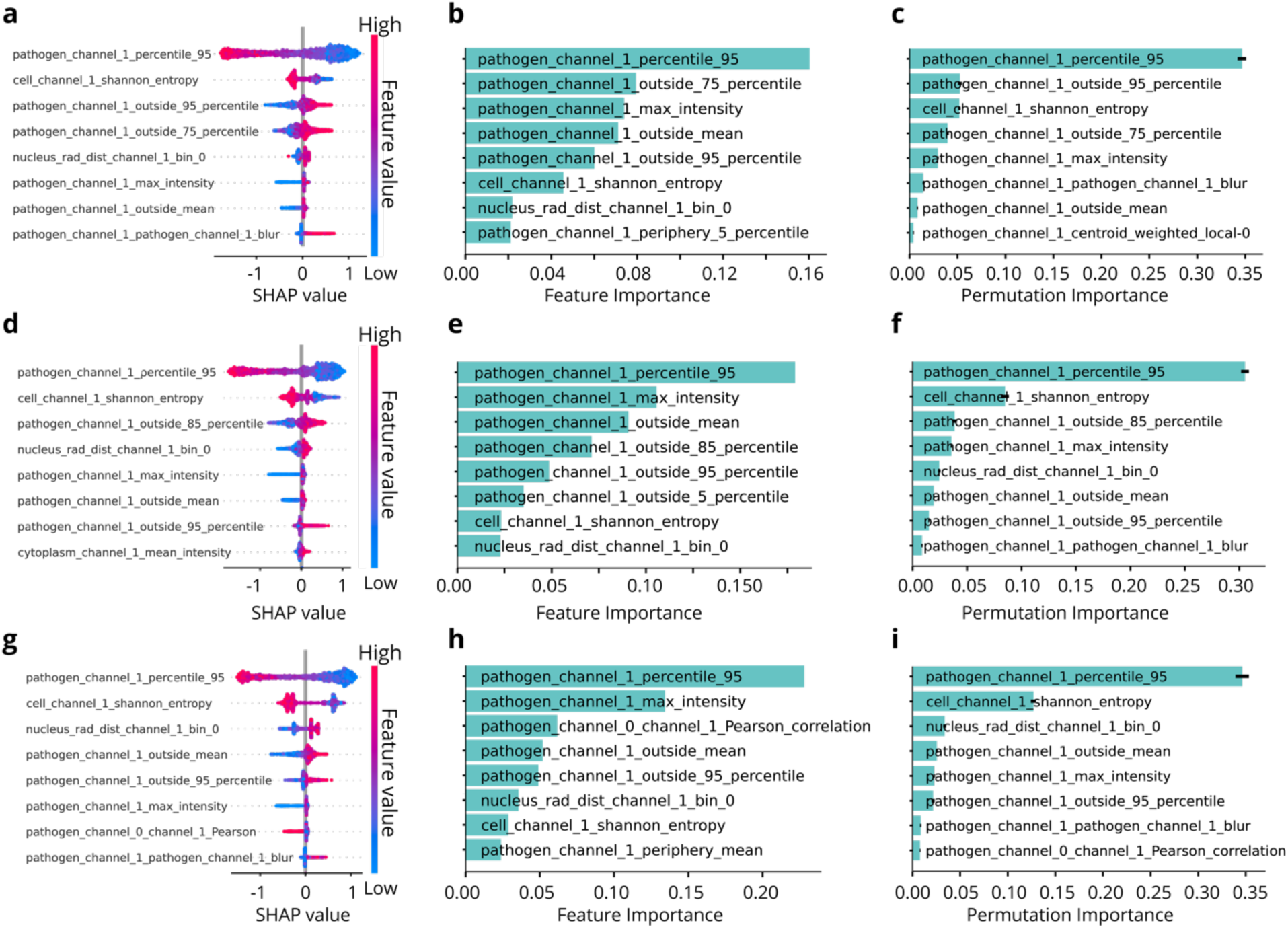
Feature attribution for XGBoost^GFP-TSG101^ models across plates. Top eight features by SHAP feature importance (***a***, ***d***, ***g***), model-derived feature importance (***b***, ***e***, ***h***), and permutation importance (***c***, ***f***, ***i***) for XGBoost^GFP-TSG101^ models trained on PC and NC cells from plate 2 (a-c), plate 3 (d-f), and plate 4 (g-i). The corresponding analysis for plate 1 is shown in Fig. 4b-d.

**Extended Data Figure 5.**
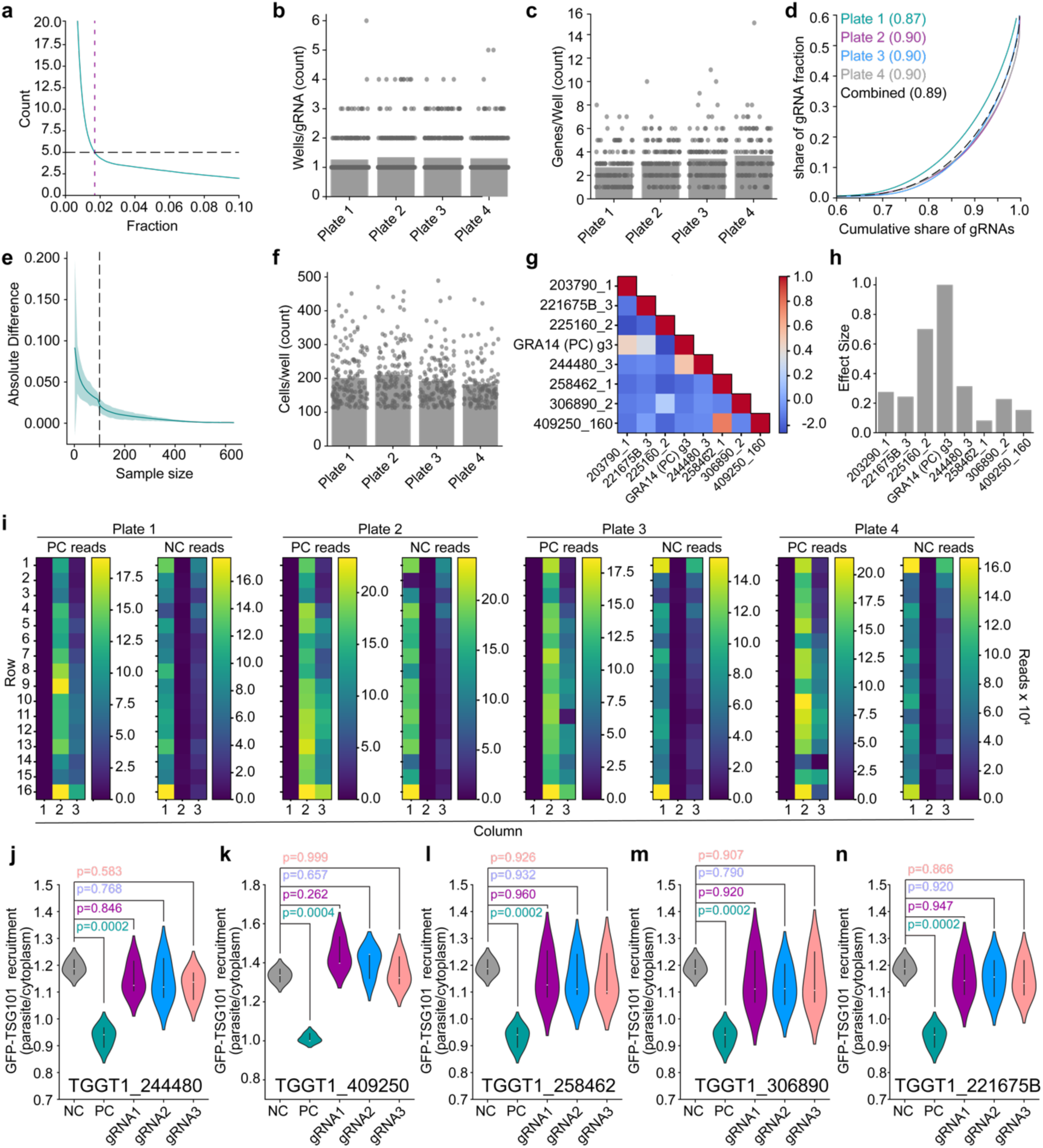
Quality control of screen variables and validation of non-confirmed candidates. ***a***, gRNA fraction threshold. Read counts for each gRNA in each well were converted to fractions. The threshold (∼0.02, dashed purple line) was set empirically as the fraction at which an average of five gRNAs per well were retained (dashed black line). ***b***, number of wells in which each gRNA was detected, by plate. ***c***, unique genes detected per well, by plate. ***d***, Lorenz curve of gRNA abundance equality in screen wells. Gini coefficients are given in brackets per plate. ***e***, cell-number threshold derivation. Cells were randomly removed from the 100 wells with the highest cell counts. The well score deviated by less than 2% from the original above 100 cells. ***f***, cells per well by plate after applying the 100-cell minimum. **g**, well co-occurrence matrix among screen hits. ***h***, estimated effect-size misattribution for screen hits assuming fixed gene-level effect sizes of 1.0 for GRA14 (TGGT1_239740_3) and 0.7 for EAF1 (TGGT1_225160_2). Panels g and h were generated with spaCR.submodules.post_regression_analysis. ***I***, read counts of the negative control (NC, TGGT1_233460_4) and positive control (PC, TGGT1_239740_1) gRNAs in control columns 1, 2, and 3 of each plate. ***j***-***n***, validation of five non-validated candidates selected for follow-up testing (TGGT1_244480, TGGT1_409250, TGGT1_258462, TGGT1_306890, TGGT1_221675B) by single-gRNA transfection of RHCas9DsRed parasites and assay of GFP-TSG101 recruitment in HeLa^GFP-TSG101^ cells at 24 h post-infection (n = 3 plates; one-way ANOVA with Tukey post-hoc test). NC and PC are the same samples shown in Fig. 5d.

**Extended Data Figure 6.**
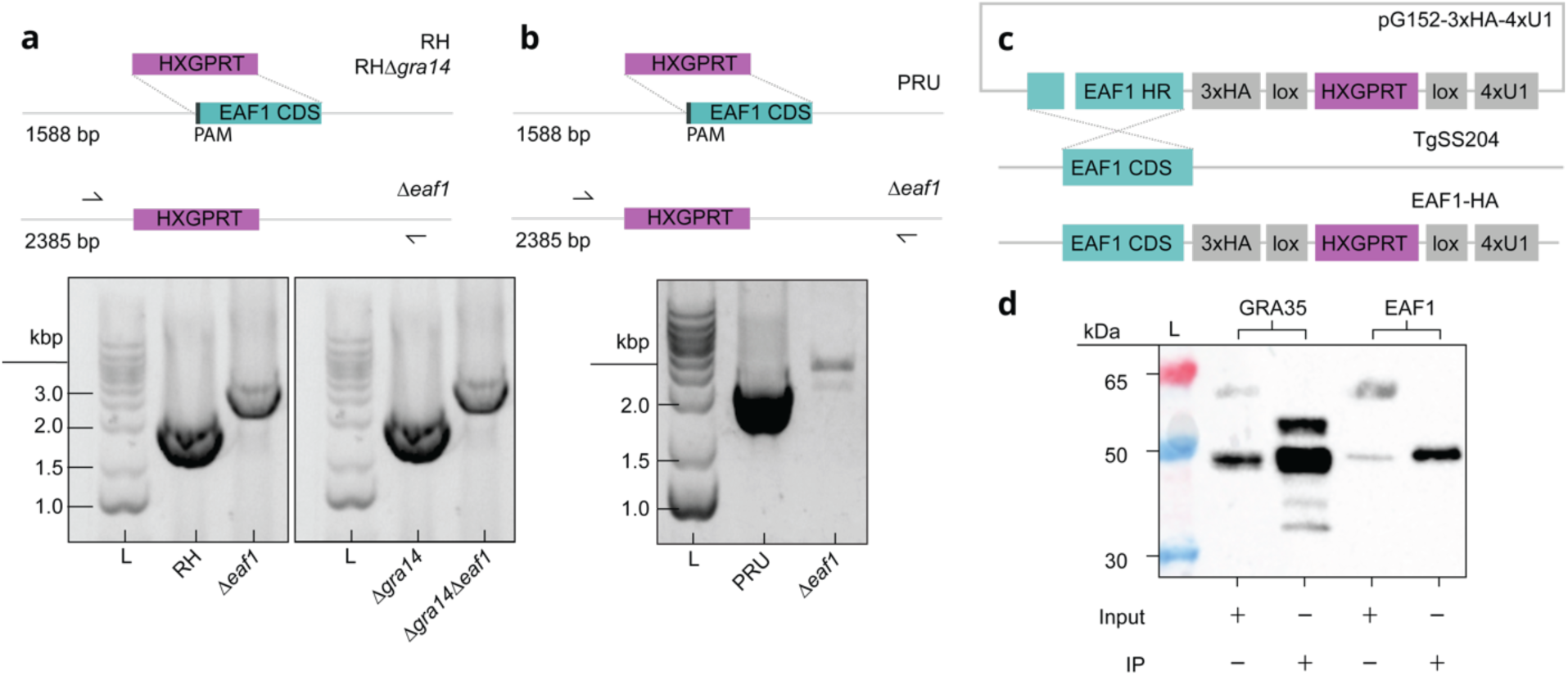
Generation of *T. gondii* strains used in this study. ***a***, Generation of RHΔ*eaf1* and RHΔ*eaf1*Δ*gra14* by integration of the HXGPRT cassette into the *EAF1* locus in RH and RHΔ*gra14* backgrounds. Diagnostic PCR confirms 5′ and 3′ integration junctions. Expected band sizes are indicated. ***b***, Generation of PruΔ*eaf1* by replacement of the *EAF1* coding sequence with HXGPRT in the PRUΔ*ku80* background. Diagnostic PCR confirms loss of the endogenous *EAF1* coding sequence and HXGPRT integration. ***c***, Strategy for the RHΔ*ku80* DiCre-based EAF1-HA tagged line. Single homologous recombination introduced a C-terminal 3xHA tag followed by a loxP-flanked HXGPRT cassette and 4xU1 terminator at the endogenous *EAF1* locus in TgSS204 parasites. Anti-HA immunoblotting confirms HA-tagged EAF1 expression (see Fig. 5g). ***d***, Representative anti-HA immunoblot of input lysates and HA immunoprecipitates (IP) from GRA35-HA control and EAF1-HA parasites. EAF1 is 45.4 kDa and GRA35 is 42.6 kDa. Both tagged proteins were detected in the corresponding input samples and were enriched after immunoprecipitation, confirming expression and recovery of the bait proteins used for IP-MS analysis.

**Extended Data Figure 7.**
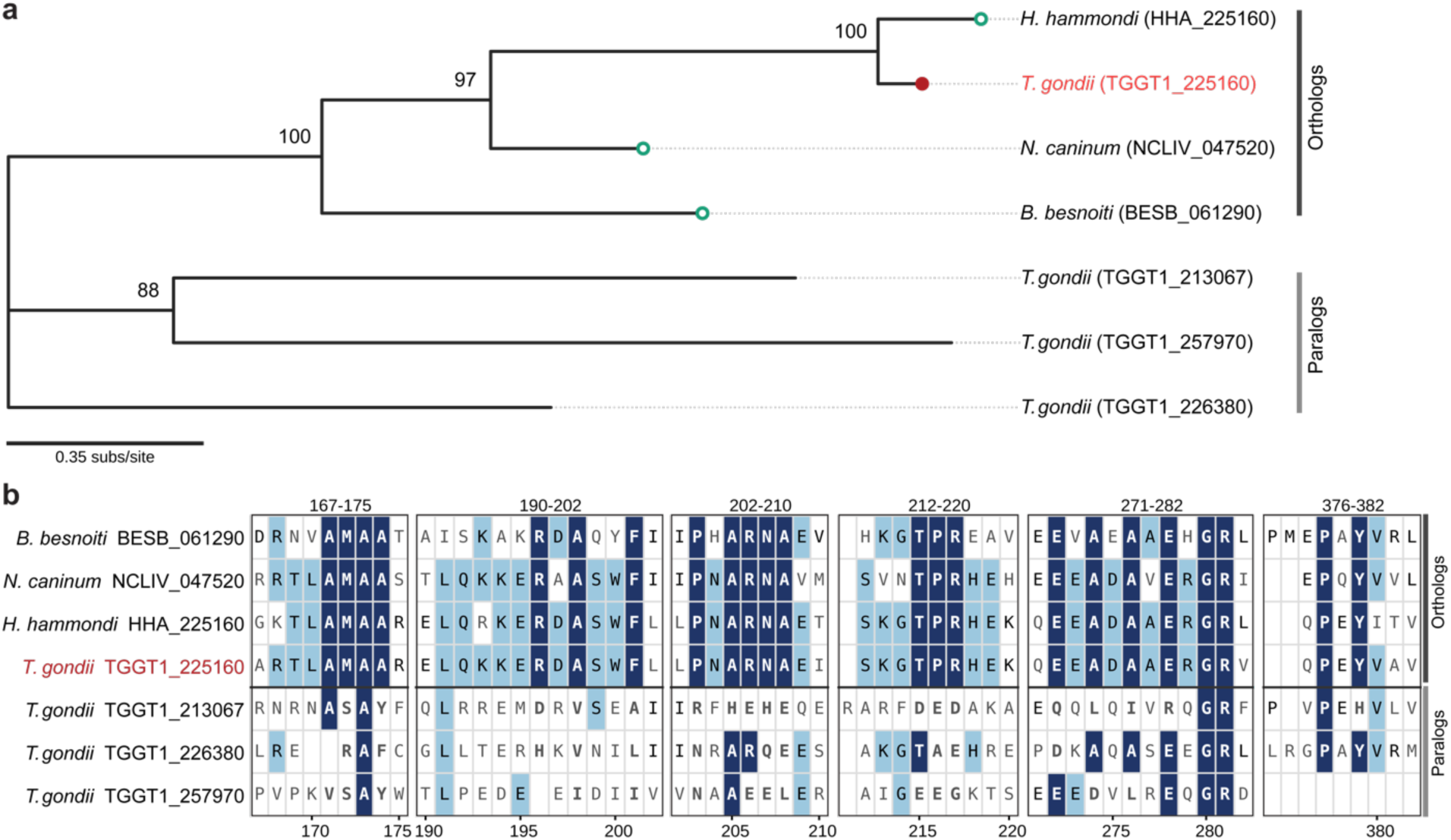
Conservation of EAF1 (TGGT1_225160) across the *Toxoplasmatinae* and within its *T. gondii* paralog family. Alignment of the EAF1 orthologues (*T. gondii* TGGT1_225160, *H. hammondi* HHA_225160, *N. caninum* NCLIV_047520, *B. besnoiti* BESB_061290), and the three *T. gondii* paralogs GRA36 (TGGT1_213067), GRA35 (TGGT1_226380) and TGGT1_257970. Residues are shaded by similarity to the consensus of the four orthologues. Dark blue marks residues identical to the orthologue consensus, light blue marks residues that are chemically similar to the consensus (conservative substitutions), and unshaded (white) cells mark residues that differ from the consensus or align to a gap. ***a***, Maximum-likelihood phylogeny. Numbers at internal nodes are local support values from the Shimodaira-Hasegawa test, shown as percentages, where 100 indicates maximal support. Circles mark the orthologues, red marks the reference TGGT1_225160. The scale bar represents 0.35 substitutions per site. ***b***, Conserved blocks in the C-terminal domain.

**Extended Data Figure 8.**
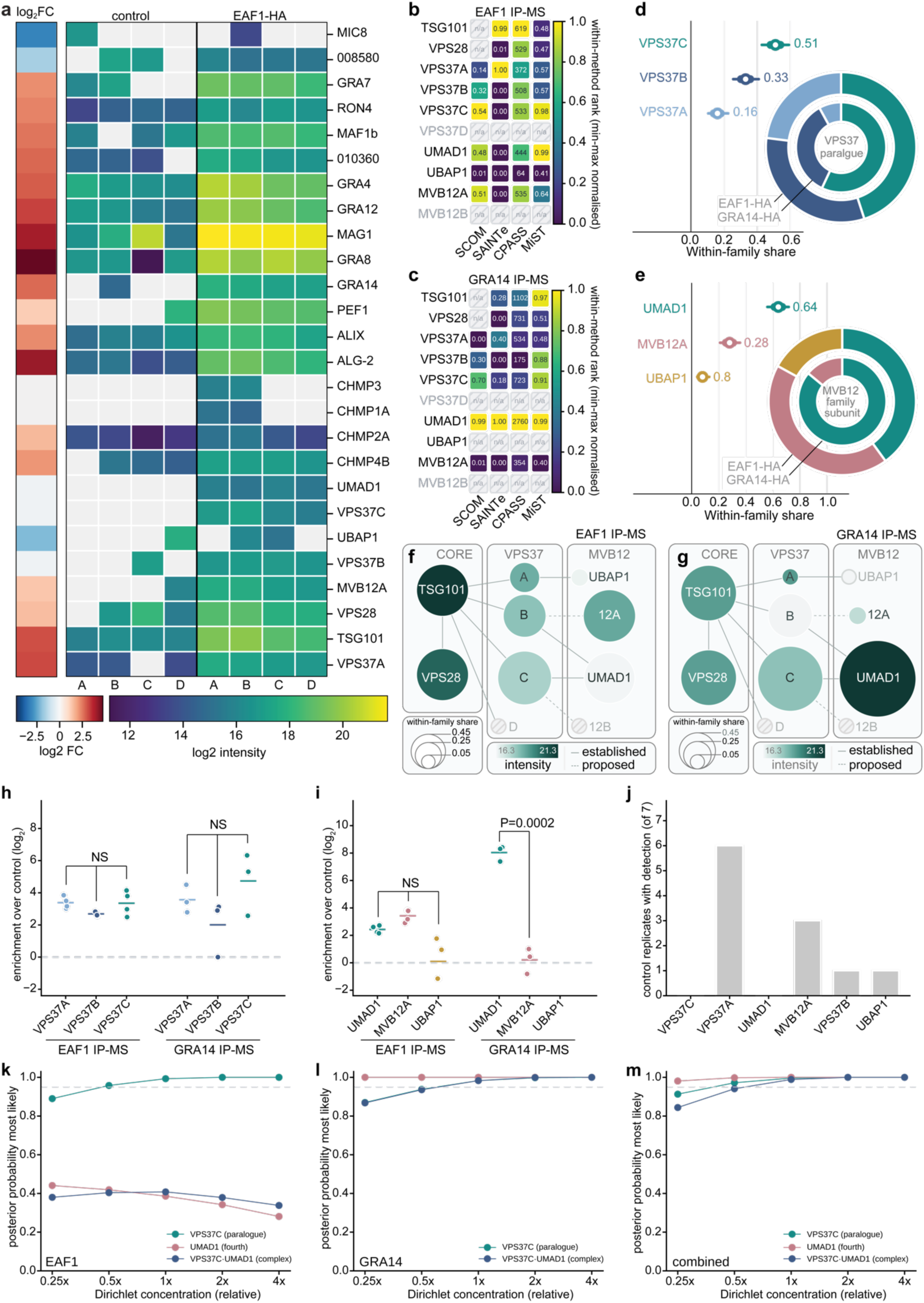
IP-MS. **a,** Anti-HA immunoprecipitation followed by LC-MS/MS on HFFs infected with EAF1-HA-tagged or untagged TgSS204 *T. gondii* (n = 4 per condition), showing all detected ESCRT associated and *T. gondii* proteins. Main panel, log_2_ intensity per replicate (A-D), missing values in grey. Left sidebar, log_2_ fold change (EAF1-HA vs untagged) on a red-white-blue scale (limma, empirical Bayes, Benjamini-Hochberg). **b**, **c**, Comparison of subunit scoring methods for the EAF1-HA (b) and GRA14-HA (c) pull-downs. Rows are ESCRT-I subunits and columns are stoichiometry-constrained occupancy model (SCOM) share, SAINTexpress (SAINTe) average probability, CompPASS WD-score (CPASS), and the adapted MiST score. Each column is min-max normalized across subunits, so colour encodes the rank within a method. Printed values are unnormalized scores. CompPASS and MiST specificity was computed against matched controls. **d**, **e**, Model-ranked VPS37 paralog (d) and MVB12-family subunit (e) recovered with EAF1 and GRA14. Forest plots rank proteins by combined posterior share (dot, median; thick bar, 50%; thin bar, 94% credible interval). Nested donut, EAF1 (outer) and GRA14 (inner). **f**, **g**, ESCRT-I interaction network for EAF1 (f) and GRA14 (g), subunits on assembly contacts, edges marking documented interactions. Node color, log_2_ bait intensity; node size, model-derived within-family relative share; undetected subunits shown as open grey markers on dashed edges. **h**, **i**, Per-replicate enrichment over control (log_2_) for the VPS37 paralogs (h) and MVB12-family subunits (i) in each dataset (points, replicates; bars, means; one-way ANOVA with Tukey’s HSD; NS = Nonsignificant). **j**, Control replicates detecting each subunit (of seven) (Fisher exact test). **k-m**, Sensitivity of model-derived relative shares to the Dirichlet concentration for EAF1 (k), GRA14 (l) and combined (m), as the concentration is scaled from one quarter to four times its value (dashed line, 0.95).

